# Neuropathy target esterase activity predicts retinopathy among *PNPLA6* disorders

**DOI:** 10.1101/2023.06.09.544373

**Authors:** James Liu, Yi He, Cara Lwin, Marina Han, Bin Guan, Amelia Naik, Chelsea Bender, Nia Moore, Laryssa A. Huryn, Yuri Sergeev, Haohua Qian, Yong Zeng, Lijin Dong, Pinghu Liu, Jingqi Lei, Carl J. Haugen, Lev Prasov, Ruifang Shi, Hélène Dollfus, Petros Aristodemou, Yannik Laich, Andrea H. Németh, John Taylor, Susan Downes, Maciej Krawczynski, Isabelle Meunier, Melissa Strassberg, Jessica Tenney, Josephine Gao, Matthew A. Shear, Anthony T. Moore, Jacque L. Duncan, Beatriz Menendez, Sarah Hull, Andrea Vincent, Carly E. Siskind, Elias I. Traboulsi, Craig Blackstone, Robert Sisk, Virginia Utz, Andrew R. Webster, Michel Michaelides, Gavin Arno, Matthis Synofzik, Robert B Hufnagel

## Abstract

Biallelic pathogenic variants in the *PNPLA6* gene cause a broad spectrum of disorders leading to gait disturbance, visual impairment, anterior hypopituitarism, and hair anomalies. *PNPLA6* encodes Neuropathy target esterase (NTE), yet the role of NTE dysfunction on affected tissues in the large spectrum of associated disease remains unclear. We present a clinical meta-analysis of a novel cohort of 23 new patients along with 95 reported individuals with *PNPLA6* variants that implicate missense variants as a driver of disease pathogenesis. Measuring esterase activity of 46 disease-associated and 20 common variants observed across *PNPLA6*-associated clinical diagnoses unambiguously reclassified 10 variants as likely pathogenic and 36 variants as pathogenic, establishing a robust functional assay for classifying *PNPLA6* variants of unknown significance. Estimating the overall NTE activity of affected individuals revealed a striking inverse relationship between NTE activity and the presence of retinopathy and endocrinopathy. This phenomenon was recaptured in vivo in an allelic mouse series, where a similar NTE threshold for retinopathy exists. Thus, *PNPLA6* disorders, previously considered allelic, are a continuous spectrum of pleiotropic phenotypes defined by an NTE genotype:activity:phenotype relationship. This relationship and the generation of a preclinical animal model pave the way for therapeutic trials, using NTE as a biomarker.

## Introduction

Biallelic pathogenic variants in the *Patatin-like phospholipase domain containing 6* (*PNPLA6*, MIM #603197) gene cause a broad spectrum of neurological disorders, including spastic paraplegia type 39 (SPG39, MIM #612020), Gordon-Holmes syndrome (GDHS, MIM #212840), Boucher-Neuhäuser syndrome (BNHS, MIM #215470), Laurence-Moon syndrome (LNMS, MIM #245800), and Oliver-McFarlane syndrome (OMCS, MIM #275400) (1–5). These disorders exhibit significant pleiotropy involving the central nervous system (CNS) and peripheral nervous system (PNS), with endocrine, ophthalmic, and hair anomalies. Patients diagnosed with SPG39 typically present with cerebellar ataxia, upper motor neuron deficits, and peripheral neuropathy. Patients with GDHS and BNHS present with hypogonadotropic hypogonadism, with BNHS additionally accompanied by chorioretinal dystrophy. Childhood-onset disorders, OMCS and LNMS, include presenting features such as chorioretinal dystrophy or anterior pituitary hormone deficiency and have trichomegaly and alopecia.

*PNPLA6* encodes Neuropathy target esterase (NTE), an endoplasmic reticulum-associated enzyme that is highly expressed in the developing human brain and eye (1). The phospholipase B activity of NTE is critical for phospholipid homeostasis and membrane trafficking (6–8). NTE is 1375 amino acids long with five protein domains: an N-terminal transmembrane domain, three cyclic nucleotide binding (CNB) domains, and a NEST domain (<u>N</u>TE-<u>est</u>erase domain) at the C-terminal end. Although the function of the CNB domains remains unknown, the NEST domain contains the catalytic residues required for phospholipid remodeling (7, 9).

Previous research has shown circumstantial evidence that NTE levels may be a key driver in disease pathogenesis. Work done by our team and others utilizing patient fibroblasts with missense and predicted loss of function (pLOF; nonsense, frameshift, and truncating) variants in *PNPLA6* lead to a reduction in NTE activity (1, 10). Furthermore, skin fibroblasts from OMCS patients with biallelic *PNPLA6* missense variants had significantly less NTE activity compared to SPG39-associated genotypes, providing evidence that different clinical subtypes might be influenced by NTE activity (1).

In affected individuals, biallelic pLOF variants are exceptionally rare. While it has been proposed that single pLOF variants are associated with retinopathy (11), truncating variants are seen in all clinical diagnoses seen in *PNPLA6* patients. Therefore, the putative relationship between the clinical subtype, genotype and NTE levels in *PNPLA6* disorders remains unclear. Such relationships, especially if recapitulated in a preclinical animal model, would open the door to the therapeutic options with NTE levels as a potential biomarker.

In this study, we examine the fundamental role of NTE activity on disease onset among *PNPLA6*-associated disorders. To investigate the genotype-phenotype correlations connected with *PNPLA6* variants, a novel cohort of 23 cases with varying clinical diagnoses contains 17 novel variants in the *PNPLA6* gene. Meta-analysis of this cohort and 95 reported individuals reveals missense variants in the enzymatic domain as a driver of disease pathogenesis due to their recurrence in specific clinical diagnoses. Using a modified NTE enzymatic assay to incorporate full-length protein analytics (12, 13), we characterize the activities of 66 missense, truncating, and common *PNPLA6* variants in vitro that was able to resolve uncertainty in all variants tested. Estimating an affected individual’s overall NTE activity by taking the average in vitro activity of their two variants demonstrates a relationship between residual NTE activity and presence of clinical subtypes. This phenomenon was recapitulated in a preclinical mouse model of the *PNPLA6* disorder spectrum, where retinopathy and mouse viability is predicated on NTE activity. Overall, these experiments uncover a novel genotype:NTE activity:phenotype relationship in *PNPLA6*-associated disorders that lay the foundation for preclinical trials using NTE activity as a novel biomarker.

## Results

### Missense variants define the PNPLA6 genotype-phenotype spectrum

To investigate the genotype-phenotype associations among *PNPLA6* disorders, we performed a clinical meta-analysis of reports of individuals with variants in the *PNPLA6* gene (1-5, 11, 14-41) along with a novel cohort of 23 patients carrying 12 reported and 17 novel variants in the *PNPLA6* gene (See Supporting Data Values xls). Through May 2023, 118 individuals have been reported (including this study) with *PNPLA6* disorders and biallelic genotypes containing 106 unique *PNPLA6* variants (Figure 1A). In total, 70/106 of variants were missense substitutions, 35/106 were predicted loss-of-function (pLOF), including splice-altering, nonsense, or frameshift variants, and 1/106 was in-frame deletion (Figure 1B). Categorizing missense and in-frame deletion variants based on protein functional domain revealed that a majority (41/71) are located within the NTE catalytic domain of the protein. Interestingly, several missense variants, such as p.Gly1129Arg, were observed to recur in association with specific clinical diagnoses. Given the prevalence of missense variants and the near absence of reported individuals with homozygous null alleles, we hypothesized that *PNPLA6* missense alleles are a key driver in NTE disease pathogenesis and determining patient phenotype.

**Figure 1.**
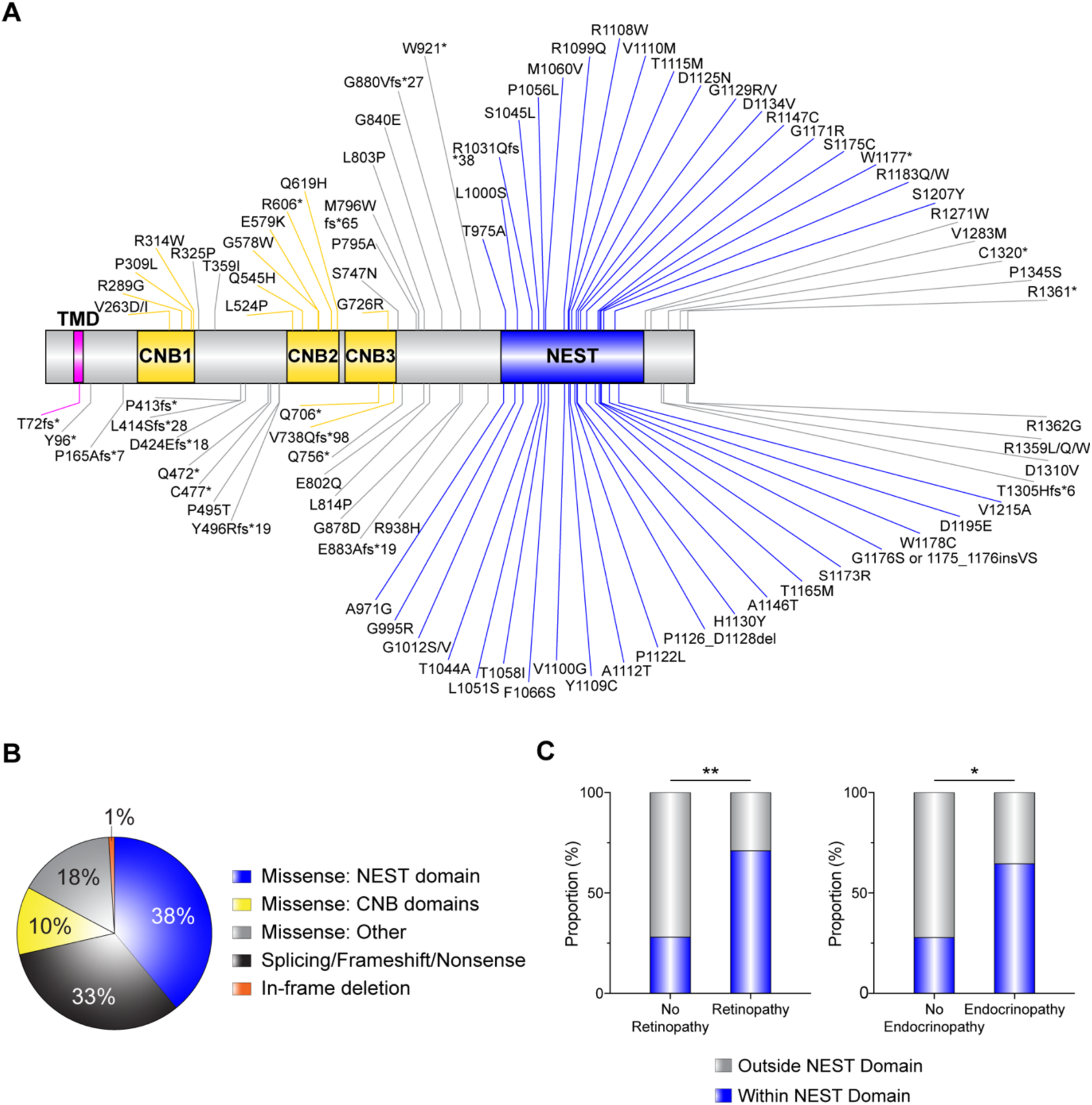
Summary of clinical meta-analysis of patients with *PNPLA6* variants. **(A)** Schematic of the protein domain architecture and the location of 93 *PNPLA6* missense, truncating, nonsense, and in-frame variants (excluding 11 splice variants, 1 duplication, and 1 deletion) known since May 2023 (NCBI reference NP_001159583.1). NTE contains a single pass transmembrane domain (aa60-80), 3 cyclic nucleotide binding (CNB) domains, and a NEST domain. **(B)** Frequency distribution of variants based on their locations on the protein and variant type. N is as follows: Missense: NEST domain: 40; Splicing/Frameshift/Nonsense: 35; Missense: CNB domains: 11; Missense: Other: 19; In-frame deletion:1. **(C)** Missense variants categorized within the NEST domain (aa964-1269) and outside the nest domain (aa1-964, aa1270-1375) were grouped by their association with/without retinopathy and endocrinopathy. (C) used a Fisher’s exact test with α=0.05.

To understand the role of missense alleles in disease severity and organ-specific phenotypes, we categorized missense variants located outside the NEST domain (aa1-963, 1270-1375) and within the NEST domain (aa964-1269) and compared their frequencies between individuals with and without retinopathy and endocrinopathy. Significantly more missense alleles located within the NEST domain are associated with disorders causing syndromic retinopathy (27/40, p=0.002) and endocrinopathy (31/40, p=0.01) compared to those not associated with retinopathy (7/40) or endocrinopathy (5/40) (Figure 1C). In contrast, this pattern was not observed in association with missense variants located outside the NEST domain. Hence, missense variants within the NEST domain are observed more frequently in individuals with retinopathy and endocrinopathy suggesting NTE function in retinal and endocrine-related tissues is more severely affected by variants in the NEST domain than outside the NEST domain.

### PNPLA6 missense variants modulate NTE activity

We hypothesized that residual enzymatic activity correlates with disease severity, including the risk of retinopathy and endocrinopathy. To evaluate the effect of missense variants on NTE enzymatic activity, we developed an enzymatic assay for testing variants in full-length PNPLA6 protein, using previous NTE assays (1, 10, 42). HEK293 suspension cells were transfected with full-length *PNPLA6* harboring individual missense mutations located throughout the gene, and enzymatic activity was measured using a colorimetric assay that directly measures NTE hydrolase activity by cleavage of phenol valerate (Figure 2A). We tested 41 missense variants, 20 common variants, 4 truncating variants, and 1 in-frame deletion variant. The NTE activity of all disease-associated missense/in-frame and truncating variants was significantly reduced compared to the wildtype control, whereas 20 common variants were not significantly different, establishing our assay’s robustness to measure NTE activity accurately.

**Figure 2.**
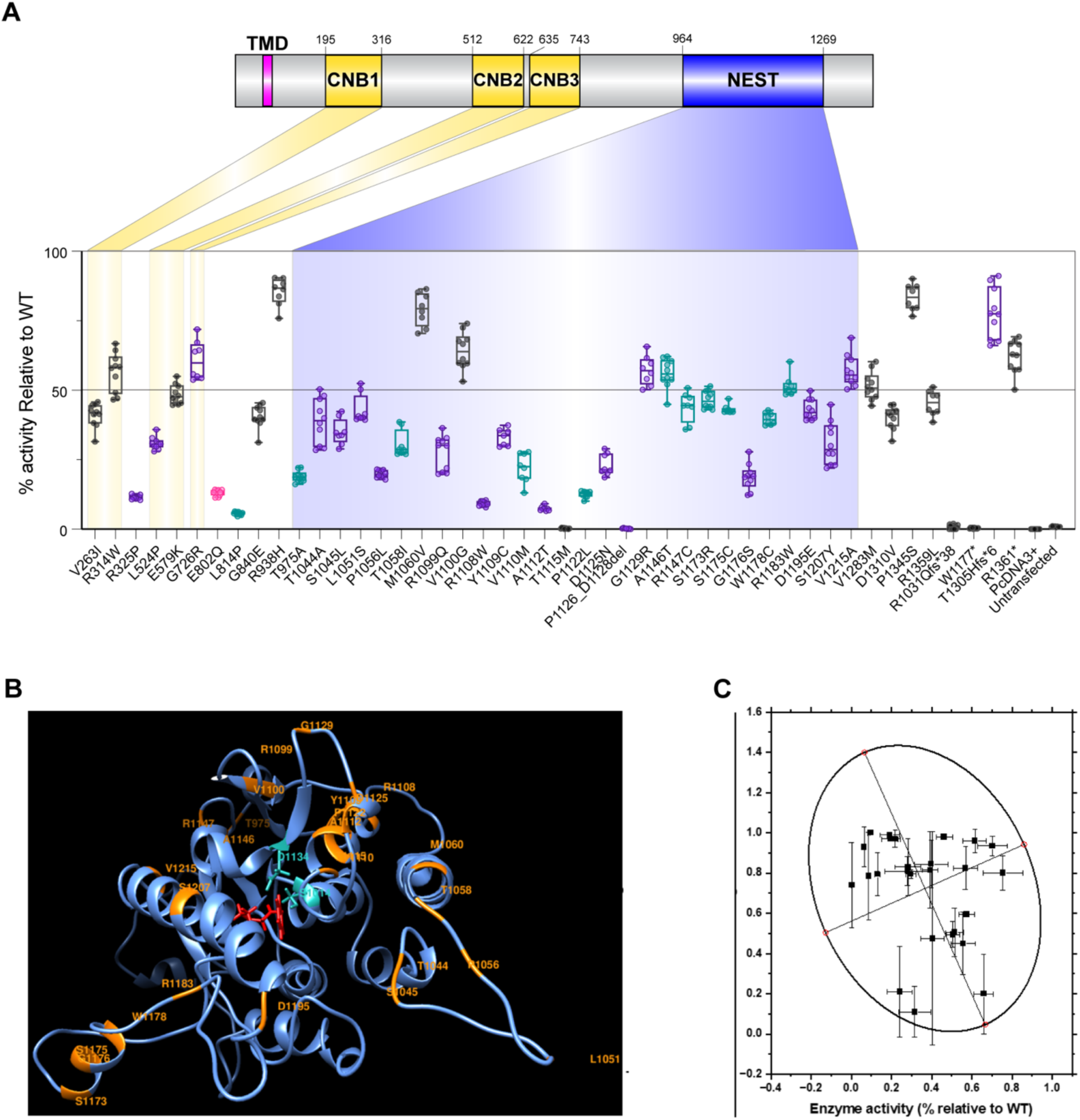
*PNPLA6* variants reduce in vitro NTE activity and increase likelihood of in silico misfolding. **(A)** Variants color-coded by the syndrome they associate with (Black=SPG39; Pink=GDHS; Teal=BNHS; Purple=OMCS/LNMS). Highlighted areas indicate the location of variants within the protein’s domains (RefSeq protein NP_001159583.1). N>7 for all samples tested. “% activity” is the activity of variants relative to WT. The activity of variants was normalized to the WT samples run in parallel. Each run had a WT, a previously tested missense variant, Arg1031Glnfs*38, and Untransfected samples. Box and Whisker plots extend from the 25th to 75th percentiles, with whiskers extending to the minimum and maximum values in the dataset. The median is the line plotted between boxes. **(B)** Shows the homology model of the human patatin domain. The ribbon structure of the patatin domain is shown in blue. Positions of 27 missense variants in the domain are labeled orange. Two residues forming a catalytic dyad, Ser1014-Asp1134, are shown in cyan. A patatin substrate, phenyl pentanoate, was docked to the active site and shown in red. **(C)** Global computational mutagenesis analysis comparing predicted unfolding fraction vs. residual NTE activity. A 95% confidence ellipse is fitted in a scatter plot. R^2^=0.7.

Truncating variants that disrupt the NEST domain (p.Arg1031Glnfs*38, p.Trp1177*) produced zero NTE hydrolase activity as expected. Interestingly, truncating variants located outside the NEST domain and near the C-terminal end of the protein (p.Thr1305Hisfs*6, p.Arg1361*) exhibited residual esterase activity, suggesting that truncation of the protein near the C-terminal end results in preserved yet decreased residual esterase activity. Three novel splice variants were clarified using a minigene splicing assay and digital droplet PCR (ddPCR). Splice variant c.1635+3G>T produced two alternative splice products, whereas c.1635+10_1635+15del produced a mixture of canonical and alternatively spliced products (Supplemental Figure 1A). This mixture of products was also seen in the splice variant c.1636-3C>A, with ddPCR quantifying the total levels of each transcript (Supplemental Figure 1, B and C).

Because phenol valerate is a synthetic substrate, we validated this observation using a biological substrate. The individual Phospholipase A1 activity on a fluorescently labeled lysophosphatidylethanolamine and A2 activity on a fluorescently labeled phosphatidylcholine of select missense and truncating variants was confirmed (see Methods), producing comparable activities to phenol valerate cleavage and to reported patient fibroblast activity assays (1, 10) (See Supporting Data Values xls). Individual variant activity and genotype (biallelic variant) activities correlated well between phenol valerate and phospholipase assays to measure NTE activity.

Next, we assessed the value of the NTE hydrolase activity assay in variant classification using the ClinGen standards for functional evidence. We applied the OddsPath equation developed by Brnich and colleagues (43), which produced an OddsPath of 20 for pathogenic variants, and 0.022 for benign variants, indicating the level of evidence as PS3 and BS3_Strong (See Supporting Data Values xls). Using the 2015 ACMG criteria (44) and the point-based system developed by Tavtigian *et al.* (45), 100% of disease-reported variants were able to be classified as likely pathogenic or pathogenic, including reclassification of 10 variants as likely pathogenic and 36 variants as pathogenic (See Supporting Data Values xls).

To further evaluate the relationship of missense variants in protein function, protein stability was assessed in 27 missense variants in the NEST domain using a homology model (Figure 2B) from available crystal structures of patatin-like proteins (see Methods). Applying global computational mutagenesis to determine the unfolding propensities of missense variants in the NEST domain revealed an inverse relationship between protein unfolding and residual NTE activity (Figure 2C, R^2^=0.7). This result indicates that catalytic function could be associated with a decrease in protein stability in the NEST domain.

### PNPLA6 disorder spectrum is determined by residual NTE activity levels

To test the hypothesis that total residual NTE activity is predictive of *PNPLA6* disorder severity, we calculated the estimated (“synthetic”) residual activity of biallelic variants and compared them to phenotypic expression. Individual genotype activity levels were determined by the average activity of the two individual variants. Next, categorization of individuals by clinical diagnosis showed a significant difference in average NTE activity levels in SPG39 patients (51%, N=12) compared to BNHS patients (28%, N=19, p=0.002 against SPG39) or OMCS/LNMS patients (28%, N=26, p=0.002 against SPG39) (Figure 3A). These results suggest that patient genotype dictates NTE activity levels in affected individuals, and that disease severity and synthetic NTE activity levels are inversely correlated.

**Figure 3.**
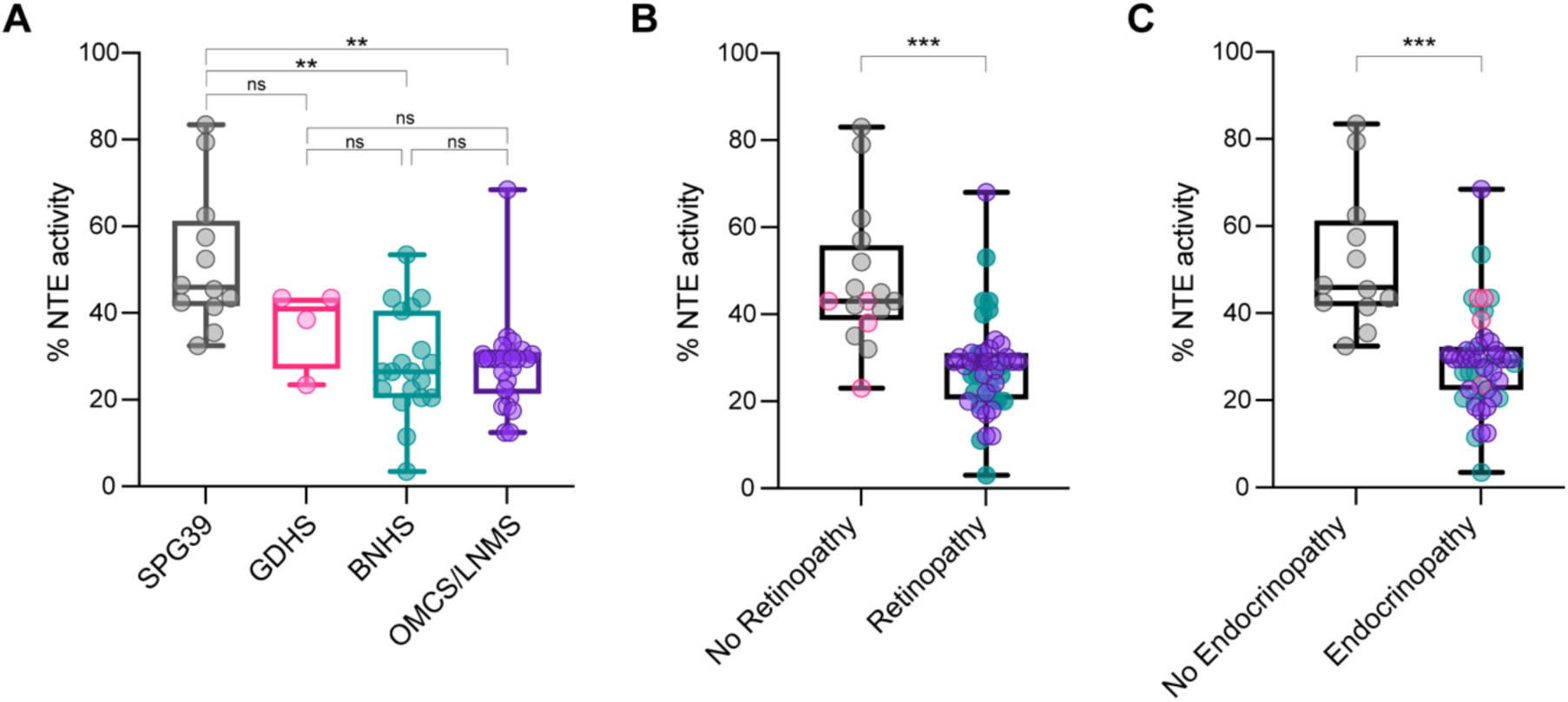
Inverse relationship between NTE PVase activity and disease severity and tissue onset. **(A)** Comparing synthetically determined patient NTE activity categorized by *PNPLA6*-associated syndromes. **(B)** Comparing synthetically determined patient NTE activity between individuals with and without retinopathy. **(C)** Comparing synthetically determined patient NTE activity between individuals with and without endocrinopathy. “% activity” is the activity of variants relative to WT. Data points for (B) and (C) correspond to the following: Black=SPG39; Pink=GDHS; Teal=BNHS; Purple=OMCS/LNMS. Box and Whisker plots extend from the 25th to 75th percentiles, with whiskers extending to the minimum and maximum values in the dataset. The median is the line plotted between boxes. (A) used a Brown-Forsythe ANOVA with post-hoc Tukey test with α=0.05. (B) and (C) used a Welch’s t-test with α=0.05. p > 0.05 = ns, p < 0.05 = *, p < 0.01 = **, p < 0.001 = ***, p < 0.0001 = ****.

As clinical diagnoses reflect the age of onset and type of affected tissues, we compared NTE activity levels in those with and without retinopathy, and in those with and without endocrinopathy. NTE activity levels in individuals with retinopathy (28%, N=45, p=0.0001) and endocrinopathy (28%, N=49, p=0.0004) were significantly lower than those in individuals without retinopathy (48%, N=16) or endocrinopathy (51%, N=12), respectively (Figure 3, B and C). Altogether, these results propose a novel genotype:NTE activity:phenotype relationship of the *PNPLA6* disorder spectrum, where lower residual activity results in a higher likelihood of more severe disease outcomes, as well as retinopathy or endocrinopathy.

### Identifying tissue-specific patterns between residual NTE activity and onset of disease

To demonstrate the relationship between NTE activity and the onset of disease, the age of tissue manifestation for unreported and reported individuals (1, 4, 28) with biallelic *PNPLA6* variants was correlated to an individual’s overall NTE activity (Supplemental Figure 3). There was no apparent relationship between the activity of NTE and the onset of retinopathy or endocrinopathy. A more detailed analysis looking at the onset of central and peripheral nervous system clinical disease, with resulting ataxia, spasticity/pyramidal tract signs, and peripheral neuropathy, did not reveal noticeable patterns either (James Liu, NIH, Bethesda, MD, USA, unpublished observations). Although there is not a discernible relationship between the onset of neurological symptoms and NTE activity, patients with overall NTE activity levels lower than 32% are highly likely to develop endocrine or ophthalmic disease findings.

### Neuropathy target esterase activity drives retinal degeneration in mice

To demonstrate that the genotype:NTE activity:phenotype relationship observed can be replicated in vivo, we developed a murine allelic series. Mice were generated with a missense variant seen exclusively in SPG39 patients (c.3088A>G, p.Met1030Val) in the homozygous state (“MV” allele), and a truncating variant “delAT” (c.3088_3089delAT, p.Met1030Valfs*2) that exhibits 0% residual activity due to its truncation prior to the enzymatic domain. We hypothesized that mice with the MV/MV genotype would have 80% NTE activity and lack retinopathy, consistent with the human phenotype in SPG39, while mice with the genotype MV/delAT (not seen in patients) and a much lower residual NTE activity, would develop retinopathy.

Mice homozygous for MV and compound heterozygous for MV/delAT were viable with typical Mendelian ratios, whereas delAT/delAT mice were not viable as predicted (Supplemental Table 1). Testing the visual function of control, MV/MV, and MV/delAT via electroretinography (ERG) over the course of 12 months revealed a significant reduction in photopic and scotopic a- and b-wave amplitudes between control and MV/delAT mice (Supplemental Figure 4). An example of this reduction in visual function can be seen in the representative scotopic and photopic traces at the max stimulus intensity (10 cd.s/m^2^ for scotopic, 100 cd.s/m^2^ for photopic) of our assay protocol at 3 months (Figure 4, A and B). Traces show a discernable reduction in both b-wave amplitudes under scotopic and photopic conditions, and smaller yet still significant differences in a-wave amplitudes. Additionally, there was no significant difference in amplitudes between control and MV/MV mice over several time points, reiterating the phenotypic observations seen in SPG39 patients who do not exhibit a retinopathy phenotype. Testing the visual acuity of mice in our allelic series at 3 months using the optomotor reflex (OMR) showed similar results, where control and MV/MV mice did not exhibit significant differences in visual acuity (+/+=0.43 cycles/degree (cpd), MV/MV=0.425 cpd p=0.80), whereas MV/delAT mice visual acuity was significantly reduced compared to controls (MV/delAT=0.378 cpd, p=0.008, Figure 4C). Overall, the reduction in ERG amplitudes and OMR provide evidence that reduction in NTE activity in a dose-dependent manner causes reduced retinal function in mice.

**Figure 4.**
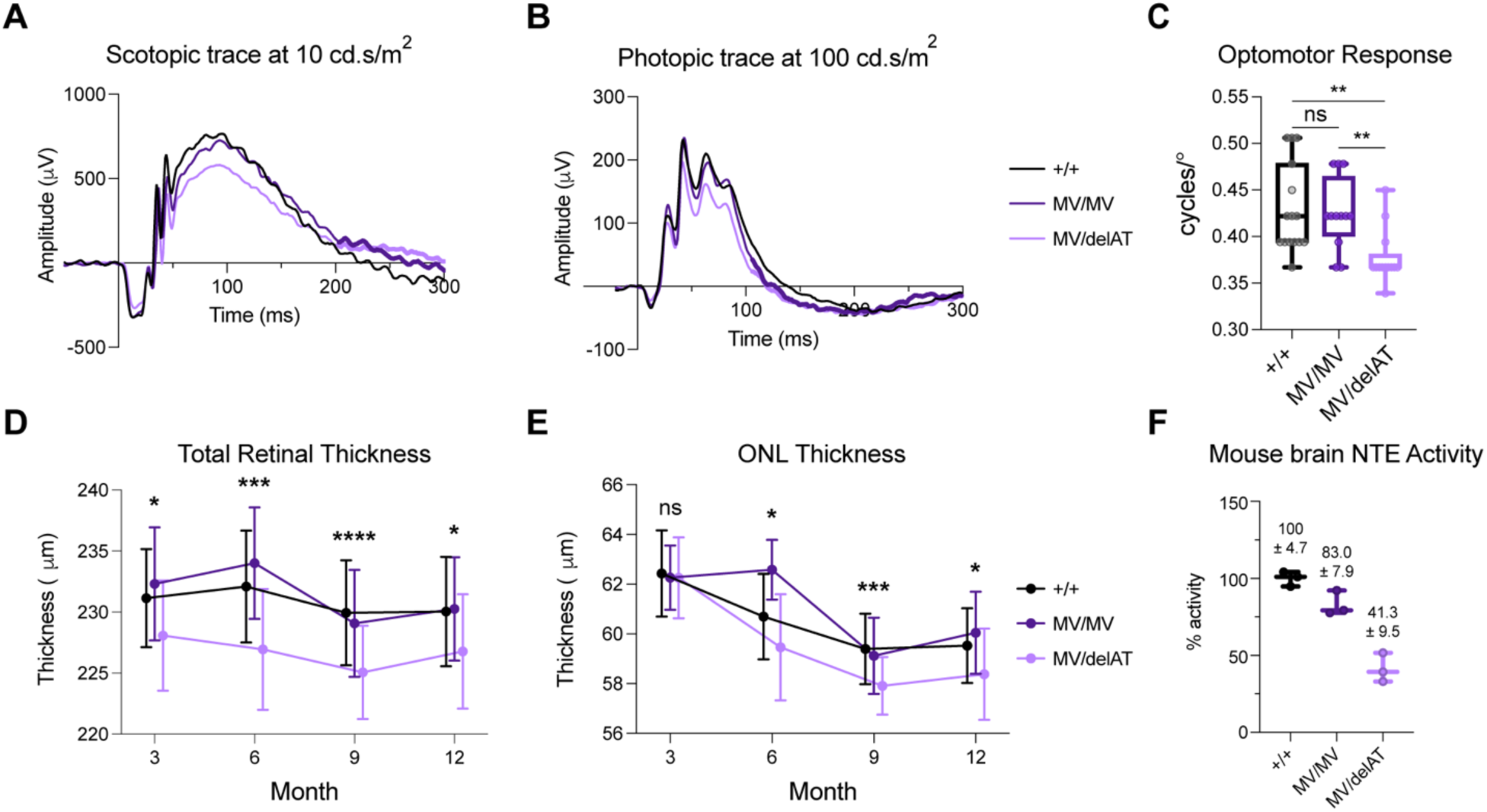
Residual NTE activity determines the loss of visual function and structure in *PNPLA6* allelic series mice in a dose-dependent manner. **(A)** Scotopic ERG trace at max stimulus intensity (10 cd.s/m^2^) for our photopic imaging protocol. **(B)** Photopic ERG trace at max stimulus intensity (100 cd.s/m^2^) for our scotopic imaging protocol. Each condition at each timepoint had an N=8-20 (see table S10 for exact N). **(C)** OMR response (+/+ N=15, MV/MV N=12, MV/delAT N=13) of allelic series mice at 3 months. **(D)** Time course of total retinal thickness of OCT images from 3-12 months. Each condition at each timepoint had an N=6-9 (see table S11 for exact N). Significant values indicate significance between Control and MV/delAT values. Control and MV/MV line were not significantly different, and MV/MV and MV/delAT were significantly different. **(E)** Time course of outer nuclear layer thickness of OCT images from 3-12 months. Each condition at each timepoint had an N=6-9 (see table S11 for exact N). Significant values indicate significance between Control and MV/delAT values. Control and MV/MV line were not significantly different, and MV/MV and MV/delAT were significantly different. **(F)** NTE activity of allelic series mice brain homogenate. N=3 biological replicates. N=3 technical replicates per biological replicate. Each timepoint took measurements from littermates, and bars in (E) and (F) are offset for cleaner presentation. All Error bars in S.D. Box and Whisker plots extend from the 25th to 75th percentiles, with whiskers extending to the minimum and maximum values in the dataset. The median is the line plotted between boxes. Statistical tests from these figures used One-way ANOVA with post-hoc Tukey test with α=0.05. p > 0.05 = ns, p < 0.05 = *, p < 0.01 = **, p < 0.001 = ***, p < 0.0001 = ****.

To examine the morphological features of the retinas in our allelic series, we obtained cross-sectional mouse retinal images by spectral domain-optical coherence tomography (SD-OCT) from 3-12 months. Using an AI-driven OCT segmentation tool, the thickness of individual retinal layers was measured (Supplemental Figure 5). The total retinal thickness measured from the retinal nerve fiber layer to the RPE showed a significant reduction in retinal thickness between control and MV/delAT mice from 3 to 12 months (Figure 4D). Similar to the visual function data, there was a significant difference in total retinal thickness at all timepoints between control and MV/delAT mice, but not between control and MV/MV mice. To identify the specific retinal layers affected by the reduction of NTE activity, individual layers (photoreceptor and RPE layer, outer nuclear layer, inner nuclear layer, inner plexiform layer, nerve fiber layer) were measured and showed intermittent differences in thickness across all timepoints between control and MV/delAT mice. Noticeably, the outer nuclear layer (ONL) was consistently thinner across all timepoints between control and MV/delAT mice (except at 3 months), and not significantly thinner between control and MV/MV mice (Figure 4E). These results reinforce the idea that NTE activity level predicts the onset of retinopathy in a dose-dependent manner.

We then measured the NTE activity of mouse brain in our murine allelic series to further validate our relative enzymatic activities measured from overexpression experiments. Using a modified NTE protocol (46) for mouse tissue, mouse brain homogenate showed similar results as observed in our in vitro results, highlighting the remarkable predictability between genotype and enzymatic activity in a dose-dependent manner (Figure 4F, Figure 5, A and C). Testing the viability of our *Pnpla6* murine allelic series revealed that less than 40% NTE activity was embryonic lethal in mice, demonstrating that a threshold of residual NTE activity is needed for proper embryonic development (Figure 5B). Evaluating the visual function and structure of viable homozygous mice in our allelic series indicated that retinal degeneration only occurs in the MV/GS mouse line, which has an overall activity level of 50% (Figure 5, D-F). This indicates that the onset of retinal degeneration occurs between 40-50% residual NTE activity, which is similar to overall NTE activity levels in patients with retinopathy. Overall, these results underpin our clinical and in vitro observations, where residual NTE activity is critical for the onset of retinopathy in individuals with *PNPLA6* biallelic variants.

**Figure 5.**
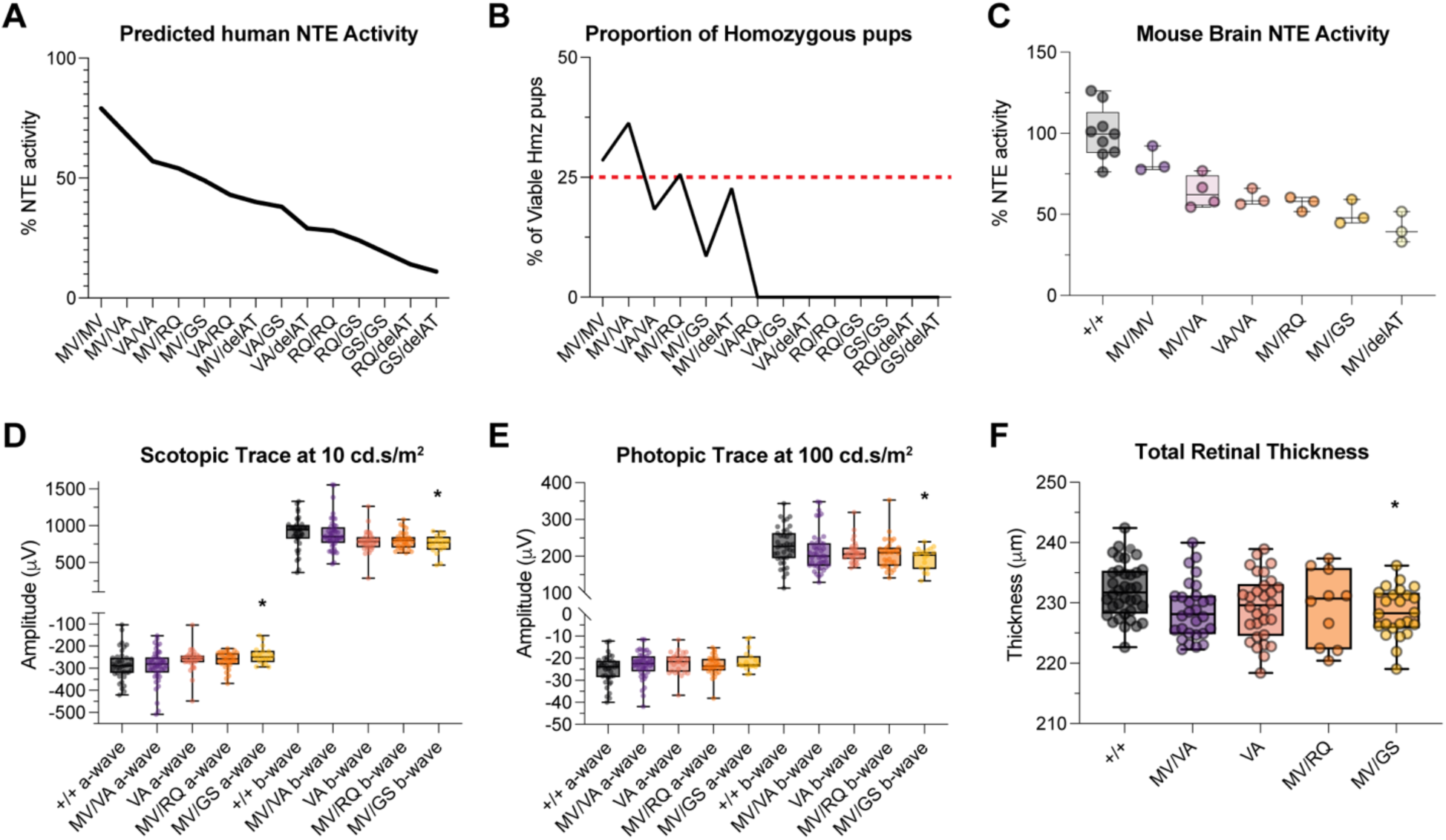
Residual NTE activity dictates the onset of retinopathy and embryonic lethality in *PNPLA6* missense allelic series mice. **(A)** Predicted overall NTE activity of allelic series mice using human in vitro variant activities (Figure 2). **(B)** The proportion of viable homozygous pups in the *PNPLA6* allelic series. The red line denotes the normal Mendelian ratio expected (25%). See table S10 for exact numbers. **(C)** NTE activity of viable homozygous allelic series mice brain homogenate. N=3-9 biological replicates. N=3 technical replicates per biological replicate. N values are found in Table S12. **(D)** Scotopic ERG wave amplitudes at a stimulus intensity of 10 cd.s/m^2^. **(E)** Photopic ERG wave amplitudes at a stimulus intensity of 100 cd.s/m^2^. **(F)** Total retinal thickness measured from SD-OCT images at 6 months. Box and Whisker plots extend from the 25th to 75th percentiles, with whiskers extending to the minimum and maximum values in the dataset. The median is the line plotted between boxes. D-F used One-way ANOVA with post-hoc Tukey test with α=0.05. p > 0.05 = ns, p < 0.05 = *, p < 0.01 = **, p < 0.001 = ***, p < 0.0001 = ****. Significant values in D-F indicate significance between Control and MV/GS values.

## Discussion

In this study, we describe a novel relationship in *PNPLA6* disorders, correlating genotype, NTE activity, and phenotype to predict the presence of retinopathy and endocrinopathy. Through a clinical meta-analysis of a novel cohort and reported patients, missense alleles were found to comprise a majority of the genotypic landscape, in particular those located within the NEST domain. This implies an important relationship between missense variants and NTE activity. Subsequent measurement of NTE activity in 66 missense, common, and truncating variants demonstrated that variants and genotypes recurrent among *PNPLA6* clinical diagnoses exhibit a broad range of residual activity. Synthetically reconstructing expected individual NTE activities based on their genotypic information produced a striking relationship between genotype, phenotype, and NTE activity, where residual NTE activity is inversely correlated with the presence of signs and symptoms of this group of disorders. We validated this relationship in a preclinical mouse model, where allelic series mice with *Pnpla6* genotypes associated with SPG39 do not develop retinopathy, while *Pnpla6* genotypes associated with OMCS develop retinopathy. Collectively, our research has established that *PNPLA6* disorders comprise a clinical spectrum in which the presence of affected tissues is determined by residual protein activity.

Clinical meta-analysis of previously individuals revealed that missense alleles directly correlate with tissue pathology. Sixty-six percent of known variants in the affected population to date are missense alleles, with 41/71 missense variants localizing within the enzymatic domain of the protein and biasing towards the more severe forms of the disease (BNHS, OMCS, LNMS). Testing the hydrolase activity of missense variants exhibited a wide range of activities, where missense variants located within the NEST domain exhibited a lower average activity compared to variants located outside the NEST domain (43% vs. 35%). Interestingly, several missense variants located near amino acids 1112-1122 had extremely low activity levels. This is possibly due to the proximity and interaction with the asparagine at position 1134, which is a crucial component of the catalytic reaction by acting as a general base and acid (9). Intriguingly, several missense variants located outside the enzymatic domain exhibited low residual activities (p.Arg325Pro, p.Leu814Pro). This could be due to disruption of the CNB domains that have unknown function thus far. Structure-function analyses of the missense variants that produce a drastic decrease in NTE activity located within and outside the NEST domain may reveal details about the abnormal catalytic mechanism and the function of the CNB domains, respectively.

Synthetically reconstructing an affected individual’s overall NTE activity revealed a striking relationship between the activity of the protein and the severity of disease that was dictated by patient genotype (Figure 3). 114/116 genotyped individuals have at least 1 missense allele, implicating missense alleles as a key driver of residual NTE activity and thus disease severity. To date, there are two patients with two frameshift variants that predict to have 0% activity (15, 41). Based on the results of this study, truncations prior to amino acid position 1177 produce 0% NTE activity, while truncations past this position produce relatively high residual esterase activity (e.g. c.4081C>T or p.Arg1361* homozygous patient having mild age of onset symptoms and diagnosis of SPG39). These findings uncover 16 additional truncating variants prior to amino acid position 1177 that are predicted to have 0% activity in the patient population. Interestingly, both patients with two truncating variants with 0% predicted NTE activity were diagnosed with GDHS. In mouse, truncation of the protein produces similar viability results, where mice with two truncating alleles (p.Met1030Valfs*2) near the start of the enzymatic domain produce non-viable pups.

Previously, *PNPLA6*-associated disorders were defined by the presence, absence, or age of onset of manifestations in specific organs or tissues (47). For example, OMCS is defined by the presence of “trichomegaly, chorioretinal dystrophy, and congenital or childhood hypopituitarism” (47). Although these disorders were previously defined, several individuals with biallelic *PNPLA6* variants do not fit the definition of a specific syndrome, such as OMCS, due to the asynchronous onset of affected tissues (e.g. hypopituitarism manifesting later) or the presence/absence of effects on other tissues (e.g. CNS symptoms in OMCS patients). Additionally, many missense variants are seen recurrently throughout the affected population and are present in multiple syndromes, lending support to the theory that variants in *PNPLA6* are involved in a set of pleiotropic disorders. Our study demonstrates that phenotypic onset, especially in the pituitary and the eye, is determined by residual NTE activity that is predicted by their genotype. A striking example of this can be seen with the missense variant p.Arg1183Trp (52% activity), which has been observed in individuals with SPG39, BNHS, and OMCS. One individual homozygous for the variant is diagnosed with SPG39 and predicts to have an overall activity of 52%, three individuals with BNS have the missense variant in *trans* with a splicing or frameshift allele and predicts to have an overall activity of 26%, and 2 individuals with OMCS have the missense variant in *trans* with another missense variant (p.Ala1112Thr) and predicts to have an overall activity of 30%. This remarkable observation validates the conclusion that *PNPLA6*-associated disorders are not distinct, but a continuous spectrum of disease (1, 4) that is influenced by the activity of NTE.

Studies in our *Pnpla6* murine allelic series are the first to model *PNPLA6*-associated retinopathy by measuring the visual function and structure of affected mice. Measuring the NTE activity of viable homozygous mice in our pre-clinical mouse model produced a remarkable activity gradient that is dictated by genotype. This is highlighted in Figure 4, where the MV/MV genotype, seen exclusively in SPG39 patients, does not exhibit signs of retinopathy in either mice or humans. In contrast, the MV/delAT genotype, although not seen in the affected population to date, halves the overall NTE activity compared to the MV/MV genotype, and has a NTE activity level similar to OMCS individuals. Evaluation of other homozygous viable mice in our allelic series indicated that less than 50% NTE activity is required for the onset of retinopathy, and greater than 40% NTE activity is required for embryonic viability. Altogether, these results correlate a decrease in NTE activity with a decrease in retinal function and structure in a dose-dependent manner, signifying the importance of NTE activity in proper retinal function in mice. Comparing the average difference in magnitude of the scotopic b-wave ERG (∼16%), scotopic a-wave (∼12%), photopic a- and b-wave (∼10%), and total retinal thickness (∼2%) measurements between control and MV/delAT mouse lines across all timepoints implies that visual function may not be affected by the total loss of cells in the retina/RPE, but disruptions to cellular connections or function. Further work characterizing the molecular mechanisms and lipid profiles that lead to retinopathy will help provide clarity on the role *Pnpla6* plays in retinal degeneration.

Testing the BMI of mice in our allelic series at 3 months yielded no significant differences between control and mutant mice, implying that hormone levels are not affected when NTE activity is reduced in homozygous viable mice (James Liu, NIH, Bethesda, MD, USA, unpublished observations). In contrast, previous papers showing that conditionally knocking out *Pnpla6* using nestin-*cre* in the CNS and PNS (6, 48) caused mice to become noticeably smaller compared to littermate controls. These findings demonstrate the limitations in using *Pnpla6* allelic mouse models to look at pituitary and growth defects, as much have a low survival rate in the setting of decreased NTE activity.

Although NTE activity can predict the occurrence of retinopathy or endocrinopathy, aggregating reported and unreported patients did not show striking patterns between the onset of clinical manifestations and activity of NTE (Supplemental Figure 3). This could potentially be due to the nature of reporting the exact onset of tissue disease phenotypes, which can be difficult. Collecting the numerical age of onsets of affected tissues in *PNPLA6*-affected individuals showed that CNS and PNS phenotypes did not show significant differences in onset across clinical categories (Supplemental Figure 6, D and E). In contrast, ophthalmic and endocrine phenotypes occurred significantly earlier in patients with OMCS/LNMS (Supplemental Figure 6, A and B). This can also be seen when categorizing patients with either a “0” or “1” truncating allele, where individuals with 1 truncating allele are more likely to experience retinopathy and endocrinopathy than individuals with two missense variants (Supplemental Figure 6, C and F). To gain a better understanding of the relationship between the activity of NTE and age of onset, accurate reporting via a prospective natural history study of individuals with *PNPLA6* variants may be imperative for further analysis of the disease gene.

With the advancement of next-generation sequencing, it is vital to properly interpret variants of unknown significance (VOUS) to validate variants identified in genes that cause Mendelian disorders. Our study has provided a new and robust method of disambiguating VOUS’s in the *PNPLA6* gene that can be adapted in research institutions and CLIA laboratories. Additionally, we were able to verify three novel splice variants using a minigene splicing assay and digital droplet PCR to detect the canonical and alternative spliced products (Supplemental Figure 1).

In conclusion, in this novel cohort of 23 patients with biallelic variants we describe 12 reported and 17 novel variants in the *PNPLA6* gene together with their clinical phenotypes. Clinical meta-analysis of 118 individuals, in vitro activity measurements of pathogenic and common variants, genotype-phenotype correlation of pathogenic and common variants, and in vivo characterization of retinal activity and structure in *Pnpla6* allelic series mice support a genotype:NTE activity:phenotype relationship that dictates the disease state with NTE activity as the common mechanism. This work outlines the importance for clinicians and geneticists to be aware of the predicted effects of NTE activity on the onset of progression of affected tissues, such as the pituitary and eye. Furthermore, NTE activity can be used as a biomarker to predict the presence or absence of affected tissues, which will be invaluable for mitigation and surveillance. Future studies will look at therapeutic interventions such as gene therapy or enzyme replacement therapy to restore relative levels of NTE activity in affected tissues to ameliorate the symptoms experienced by individuals affected by *PNPLA6* variants.

## Materials and Methods

### Sanger Sequencing

*PNPLA6* genomic DNA coding regions and flanking intronic sequence were amplified by PCR. PCR amplification was performed with the Applied Biosystems (ABI) BigDye Direct kit PCR master mix or NEB OneTaq standard buffer. PCR amplicons were used as sequencing templates as recommended in the BigDye Direct kit manual. Sequencing amplification was performed using the ABI BigDye Direct kit sequencing mix. Reactions were carried out in a PCR thermal cycler and purified with Edge Biosystems Performa DTR gel filtration cartridges. Sequencing products (3 μl) were resuspended in ABI Hi-Di Formamide (10 μl). Sequencing was performed with a Genetic Analyzer and analyzed with the Mutation Surveyor (5.1.0, SoftGenetics) software.

### PNPLA6 Sequencing primers with M13 tails

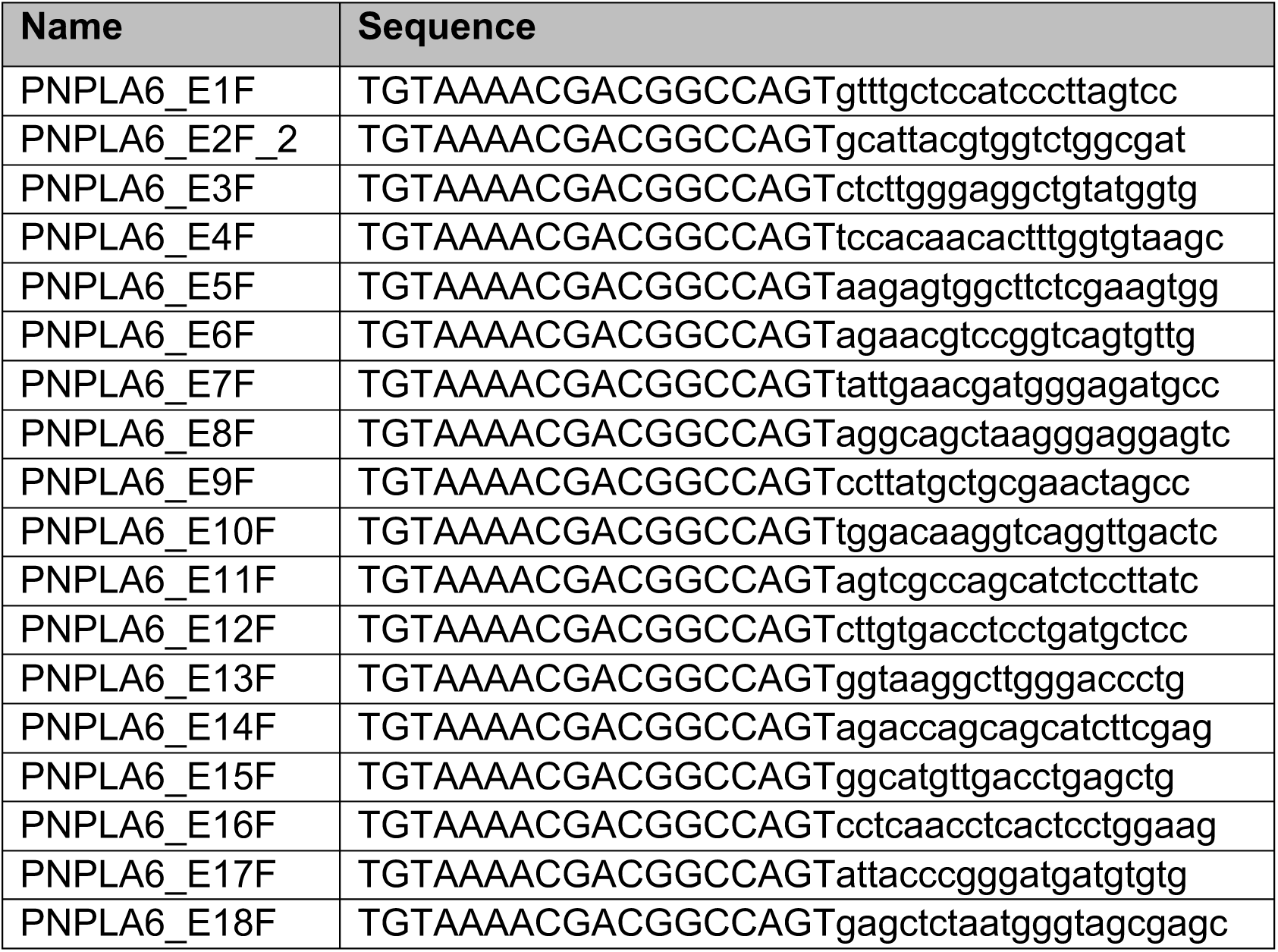
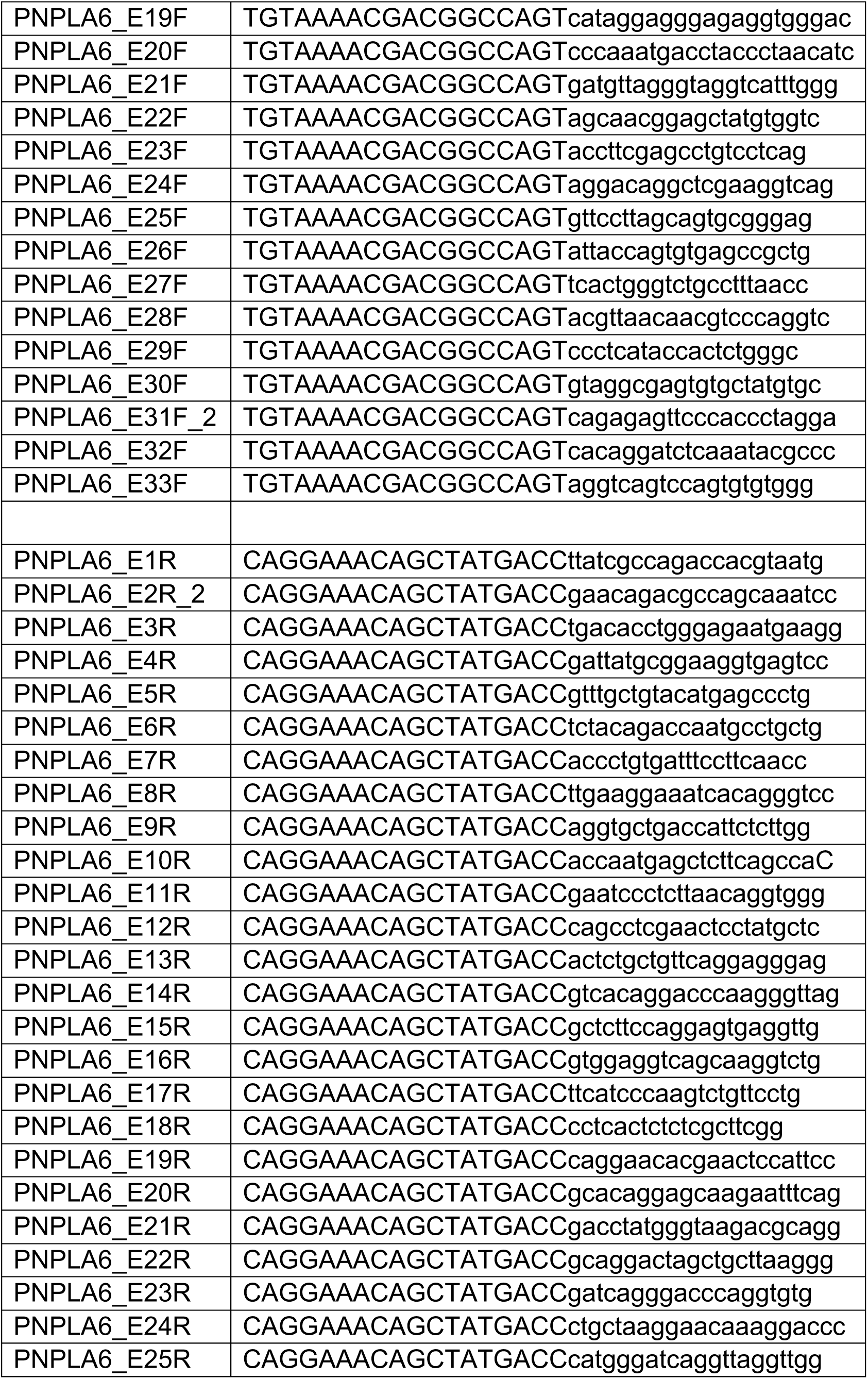
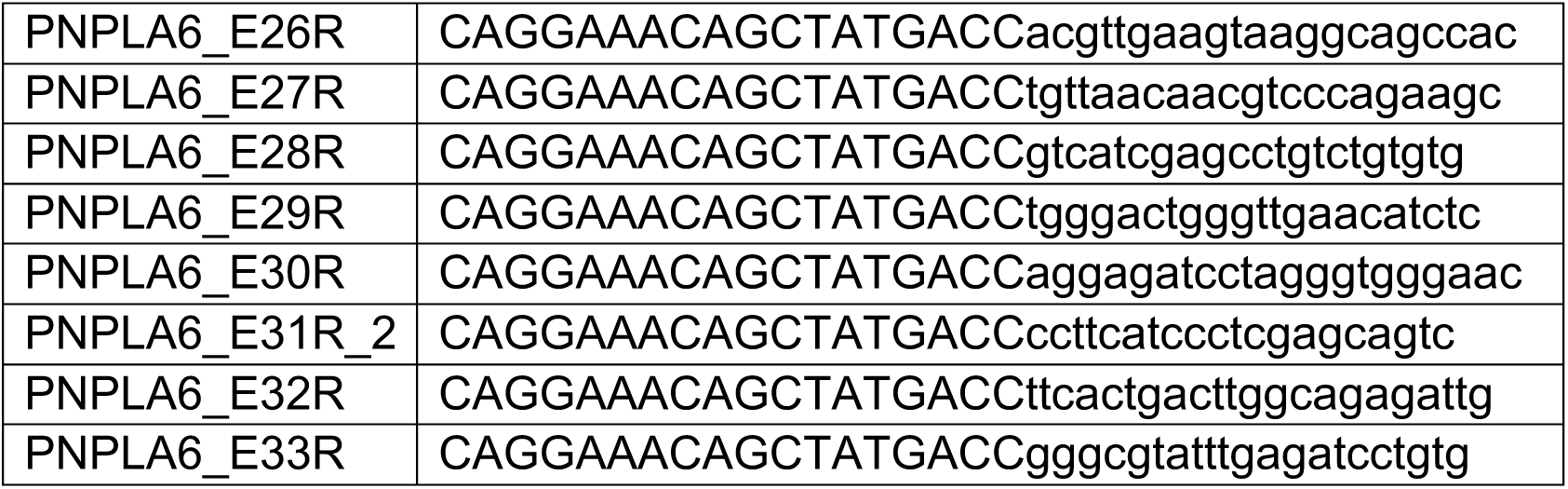

### Minigene Assay

The plasmid RHCglo used for minigene assay was a gift from Thomas Cooper (Addgene plasmid #80169 ; http://n2t.net/addgene:80169 ; RRID:Addgene_80169) (49). A DNA fragment including the *PNPLA6* exons 16, 17, and 18, plus 402 bp upstream and 307 bp downstream was synthesized according to the human genome GRCh37 and cloned into RHCglo to make the wildtype RHCglo-PNPLA6 vector, which was then used as a template to make the c.1636-3C>A, c.1635+3G>T, and c.1635+10_1635+15del mutant vector by mutagenesis (LifeSct). The wildtype and mutant RHCglo plasmids were transfected into 293FT cells using the Lipofectamine 2000 reagent (Cat# 11668019, ThermoFisher) using a suspension transfection method as described previously (PMID: 27745835). RNA was prepared from cells harvested one day post-transfection (RNeasy mini kit, Qiagen) and used for cDNA synthesis (iScript cDNA synthesis kit, Bio-Rad). M13-tagged PCR primers located in exons in the RHCglo vector were used for PCR amplification of the cDNA, RSV5U_M13F, GTAAAACGACGGCCAGTCATTCACCACATTGGTGTGC, and RTRHC_M13R, CAGGAAACAGCTATGACCGCTTTGCAGCAACAGTAACCAG. The PCR products were then subjected to Sanger sequencing using the BigDye Direct Cycle sequencing kit followed by analysis using a SeqStudio genetic analyzer (ThermoFisher).

The QX200 ddPCR EvaGreen Supermix (Cat# 186-4034, Bio-Rad) was used for RT-ddPCR using the QX200 Droplet Digital PCR System (Bio-Rad). Alternative transcript cDNA was amplified using RSV5U_M13F and PNPLA6_Intron15-M13R, CAGGAAACAGCTATGACCgatgtgatgaacacagaacc. RSV5U_M13F and PNPLA6_cDNAE16R, CTCCGCCTTGTCGATCATGC were used to determine the total amount of transcript. The canonical transcript was found by subtracting the alternative transcript from the total. GAPDH primers were TTGGTATCGTGGAAGGACTC and ACAGTCTTCTGGGTGGCAGT.

### Meta-analysis of Previously Published Patient Data

Published clinical data of patients with *PNPLA6* variants were found by the following search terms in Pubmed: “PNPLA6”, “Neuropathy target esterase”, “Oliver McFarlane Syndrome”, “Boucher Neuhäuser Syndrome”, “Gordon Holmes Syndrome”, “Spastic Paraplegia type 39”, “Ataxia”. Available patient data were recorded, including genotype, phenotype (with tissue age of onsets if available), age, sex, and diagnosis. The last search was done on May 2023.

All meta-analyses were done by patient pedigree (except for tissue age of onset analysis). Analysis of the age of onsets (AOO) in affected tissues was done with patients with available numerical AOO information. Clinical definitions for each tissue system contained at least one of the following key words:

| **Tissue system** | **Definition** |
| --- | --- |
| Central Nervous System | • White matter lesions<br>• Cerebellar atrophy<br>• Ataxia, spinocerebellar ataxia<br>• Hyperreflexia<br>• Hypertonia |
| Peripheral Nervous system | • Peripheral neuropathy<br>• Muscle numbness, tingling, and/or weakness<br>• Muscular atrophy |
| Endocrine | • Hypogonadism/Amenorrhea/Micropenis<br>• Short stature<br>• Hypopituitarism<br>• GH, LH/FSH deficiency<br>• Hypothyroidism |
| Ophthalmic | • Choroidal atrophy<br>• Abnormal retinal pigmentation<br>• Retinal degeneration<br>• Nyctalopia<br>• Vision loss |
| Hair Anomalies | • Trichomegaly<br>• Alopecia<br>• Bushy Eyebrows |

*PNPLA6* variant location were defined by RefSeq transcript NM_001166111.2. *PNPLA6* amino acid and domain locations were defined by RefSeq protein NP_001159583.1.

“0” truncating patients were defined by homozygous or compound heterozygous patients with two missense variants only. “1” truncating patients were defined by compound heterozygous patients with one missense allele, and one splice, frameshift, or indel variant.

### Variant classification

Variants were classified as benign (B), likely benign (LB), likely pathogenic (LP), and pathogenic (P) according to the 2015 ACMG/AMP guidelines (44) and point based system developed by Tavitgan *et al.* (45). Healthy population frequency data was obtained from gnomAD (v2.1) (“gnomAD browser, PNPLA6,” 2022). PS3_Strong and BS3_Strong were applied using the odds of pathogenicity (OddsPath) equations from Brnich *et al.* (43). PS4_Strong was applied to variants that had an Odds ratio >5, did not include 1, and had an adjusted α-value threshold less than 0.0002 (Bonferroni correction of 0.05/206) based on the total number of variants in the cohort. PM1 was applied to variants located in the NEST domain using RefSeq protein NP_001159583.1 (AA 964-1269). PM2 was applied to variants that were not present in gnomAD (n=0). PM3 was applied to variants that were in *trans* with variants that were previously classified as pathogenic through ClinVar or this study. PM5 was applied to variants that had a novel missense change at an amino acid residue that had been previously classified as pathogenic. PP1 was applied to variants that co-segregated in multiple affected family members with biallelic *PNPLA6* variants. BA1 was applied to benign variants that had an allele frequency >5% in gnomAD. BS2 was applied to benign variants that were classified as “gnomAD homozygotes” in gnomAD database.

### Chemicals for NTE activity assay

Phenyl Valerate was purchased and synthesized by AccuStandard (Cat# P-734N). N,N’-Diisopropylphosphorodiamidic fluoride (mipafox, MIP) was purchased and synthesized by Combi block (Cat# QZ-2872). Paraoxon (PO) was purchased and synthesized by Sigma–Aldrich (Cat# 36186-100MG). All other chemicals were reagent grade or the highest grade commercially available. All aqueous solutions involved in the enzyme assay were prepared with UltraPure DNase/RNase-Free Distilled Water. Since MIP and PO are considered neurotoxic, these chemicals were handled in the fume hood. All items in contact with MIP and PO were decontaminated by soaking in 1M NaOH overnight.

### Plasmid synthesis, mutagenesis, and purification

The plasmid pcDNA6.1/V5-His B (Thermofisher) was used as the template to subclone in the full-length *PNPLA6* cDNA (from RefSeq transcript NM_001166111.1), along with an Nluc tag on the 3’ end. This template was used to mutagenize base pairs to produce the desired patient-specific missense mutations. All plasmids were verified by DNA sequencing. Subcloning, mutagenesis, and DNA sequencing were provided by LifeSCT. Each plasmid was amplified and purified using PureLink HiPure Plasmid Filter Maxiprep Kit (Cat# K210016, Thermofisher).

### Protein expression and Cell lysate preparation

The wildtype, missense variants, predicted loss of function variants, and mock plasmids were transfected into HEK293 suspension cells derived from tsA201 (Millipore Sigma, 96121229-1VL) using Polyethylenimine (Polysciences, Cat# 24765) at a 4:1 PEI:DNA mass ratio and hybridoma SFM (Cat# 12045076, Thermofisher). An empty vector pcDNA3.1+ (Cat# V79020, Thermofisher) was used as a transfection control. The transfected cell pellet was harvested 48 hours later by centrifuging at 1000 x *g* at 4°C for 10 minutes and stored at -80 °C until needed. Thawed cell pellet was washed once with 50 mM Tris-HCl, 0.20 mM EDTA, pH 8 (enzyme assay) buffer, and recentrifuged at 1000 x *g* at 4°C for 10 minutes. After discarding the supernatant, the cell pellet was then resuspended in enzyme assay buffer again and sonicated at 3-second intervals for 2 minutes at 50% amplitude (Cat# FB120220, Fisher Scientific). The cell lysate was centrifuged at 1000 x *g* at 4°C for 10 minutes, and the resulting supernatant was used for subsequent analyses.

### in vitro NTE-specific activity assay

NTE enzymatic activity assay was performed using previously described colorimetric assays (12, 13). Briefly, cell lysate was incubated in 40 μM paraoxon and 50 μM mipafox for 20 min. Phenyl valerate (0.5mM, 0.03% w/v Tx-100) was added and incubated for 20 min. Thereafter, 1.23 mM 4-Aminoantipyrine (33.25 mg/ml SDS) was added to stop the reaction. 12.1 mM Potassium Ferricyanide was used for colorimetric determination. NTE activity is defined as the difference in phenyl valerate hydrolysis activity inhibited by paraoxon (inhibits background esterase activity) and paraoxon + mipafox (inhibits NTE in addition to background esterase activity). Endpoint absorbance was measured at 486 nm using a Synergy 2 microplate reader (Biotek). Specific NTE protein concentration was determined by SDS PAGE and Coomassie using previously defined methods (50). Li-Cor Odyssey DLx and Image Studio Lite (Li-Cor) were used to image and quantify the intensity of the bands. Band intensity was converted to a protein concentration using a BSA standard curve ranging from 0.25-1mg/ml on the same gel. The final protein concentration was determined by taking the average concentration of at least three technical replicates. Endpoint absorbance was normalized to specific NTE concentration. This assay defines one unit of NTE-specific activity as 1 μmol phenol produced per minute per milligram protein.

### Phospholipase A_1_ and A_2_ activity assay

Phospholipase A_1_ and A_2_ activity were determined using commercial assays (Cat# E10217 and E10219, Thermofisher). The protocol for determining activity was based on the manufacturer’s instructions. Cell lysate from select variants, WT plasmid, untransfected, and PLA_1_ or PLA_2_ purified enzyme (provided by Thermofisher) was used in the assay. The activity was calculated by taking the velocity at a set time point (20 min) and normalizing to protein concentration via Coomassie as stated previously. The buffer used 50 mM Tris-HCl, 0.20 mM EDTA, pH 8 (enzyme assay) buffer for all reactions. Fluorescence emission (528, 590 nm) was measured over the course of 3 hours using a Synergy 2 microplate reader (Biotek) with excitation at 485 nm.

### Molecular Modeling and Simulation

The human PNPLA6 amino acid sequence (acc# Q8IY17) was obtained from the UniProt database (https://www.uniprot.org). Residues 968 through 1286 were extracted from the *PNPLA6* sequence and were used to model the human patatin domain. The following files were downloaded from the RCSB website (https://www.rcsb.org/): Pseudomonas aeruginosa patatin-like protein structure, PLPD (5fya); human cytosolic phospholipase A_2_ dimer(1cjy); Solanum cardiophillum SeMet patatin (1oxw). Both structures, 1cjy, and 1oxw function as lipid acyl hydrolases with Ser-Asp catalytic dyad in an active site. The homology model of the human patatin dimer was generated using these proteins as structural templates in the molecular graphics, -modeling, and - simulation program YASARA (http://www.yasara.org/). The homology model of the human patatin dimer was equilibrated by 10 ns molecular dynamics in water using YASARA’s ‘run. mcr’ macro. Ion concentration was added as a mass fraction with 0.9% NaCl. The simulation temperature was set to 298 K with a water density of 0.997 g/mL. The cell size extended to 10 Å beyond each side of the protein in the shape of a cube with dimensions 119.7Å x 119.7Å x 119.7Å. Each simulation was run in YASARA using an AMBER14 forcefield, with a timestep of 1.0 fs. Simulation snapshots were outputted for every 0.1 ns.

### Global computational mutagenesis

Global mutagenesis was conducted on a model of the human patatin dimer, and each mutant was characterized by a thermodynamic change in Gibbs free energy (DDG) and by the fraction of protein in the unfolded state for each subunit (A and B) of the dimer (51, 52). Gibbs’s free energy changes were calculated using the semi-empirical method FoldX (53). The fraction of protein in the unfolded state was standardized on a 0–1 scale as the unfolding fraction u (51). The program also outputted a foldability parameter to show critical residues in protein folding (54). This parameter is a sum of severity-weighted unfolding propensities for the 19 mutations generated at a specific residue. Residues with the highest foldability were considered critical for protein folding (55, 56). Finally, to correlate the result with the enzymatic activity change caused by the mutation the unfolding parameters of subunits A and B were averaged, and standard deviations were calculated.

### Molecular docking

The structure of phenyl pentanoate was obtained from the PubChem database (https://pubchem.ncbi.nlm.nih.gov/compound/29950) of chemical structures and it was docked to the area of the catalytic dyad Ser1014-Asp1134 located in the proposed active site of human patatin homology model. The docking was performed using the Autodock tool in Yasara. For this purpose, the receptor box was centered around the catalytic dyad, and the box was set to 10 Å within the centroid.

### Generation of allelic series mice

*Pnpla6* missense variants were generated using CRISPR/Cas9 mediated homologous recombination as previously described (57). The truncating variant p.Met1030Valfs*2 (c.3088_3089delAT) used in this study was a by-product of indel mutagenesis from the production of c.3088A>G missense mutation. C57BL/6 zygotes were microinjected with a mixture of SpCas9 protein, a gRNA specific for each target and a mutagenic oligo that carries the intended knock-in mutation and short homology arms. Injected zygotes were transferred into the oviducts of female mice, and newborn F0 mice were screened for the intended knock-in mutations by PCR and sanger-sequencing spanning the knock-in or deletion area. F0 founders carrying the desired mutations are genetic mosaics and were backcrossed to C57BL/6J mice for germline transmission of each mutant allele.

Mice backcrossed greater than F4 generation were used for retinal function and structure analysis. Animals used for function and structure assays had evenly distributed males and females. Below are the sgRNA and mutagenic oligos used for this study (Targeted DNA base pair highlighted in red).

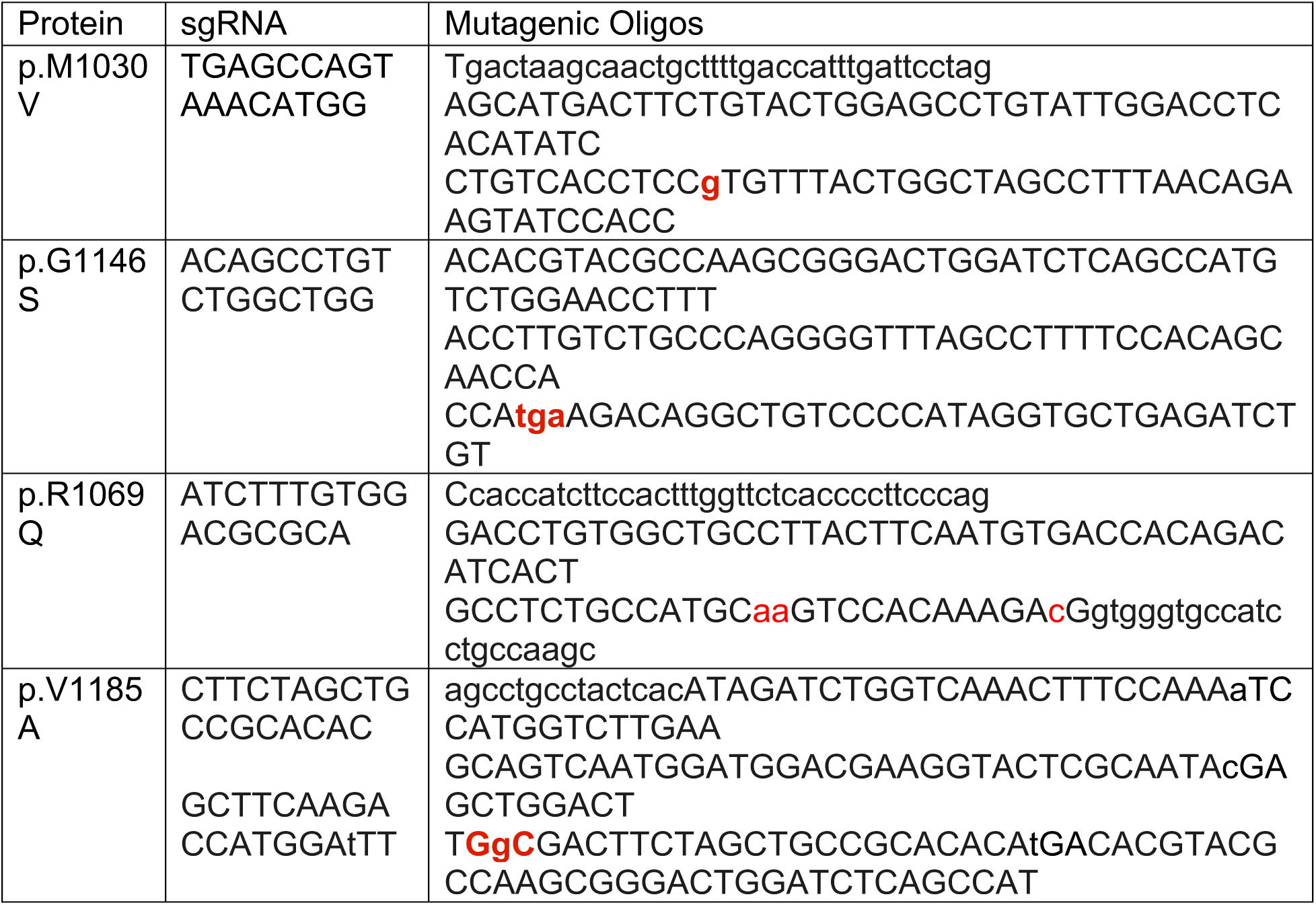

### Mouse brain preparation and activity assay

Allelic series mice at 3 and 12 months were sacrificed, and the brain was resected for activity assay analysis. Protocol was adapted from Quistad *et al.* (46). Brain (stored at - 80°C) was homogenized at 20% w/v in 50mM Tric HCl containing 0.2mM EDTA (pH 8.0). Activity assay used the supernatant fraction after centrifugation at 700x*g* for 10 min. Brain lysate was frozen down and then subsequently used for activity analysis. Activity assay of mouse brain homogenate (20 μl) was incubated in 40 μM Paraoxon and 500 μM Mipafox for 20 min. 1.5 mM Phenyl valerate (0.03% w/v Tx-100) was added and incubated for 20 min. Endpoint absorbance was measured at 486 nm using a Synergy 2 microplate reader (Biotek). Bradford assay (Cat# 5000201, Biorad) was used to determine the total protein concentration of each sample. Endpoint absorbance was normalized to total protein concentration. This assay defines one unit of NTE-specific activity as 1 nmol phenol produced per minute per milligram protein.

### Electroretinogram Recording

Electroretinogram (ERG) recordings used the Espione E3 system (Diagnosys LLC.). Mice were dark-adapted overnight (>12 hr) and anesthetized with an intraperitoneal injection of ketamine (100 mg/kg) and xylazine (6 mg/kg). All procedures were done under dim red-light conditions. Pupils were dilated with 0.5% tropicamide and 2.5% phenylephrine, and anesthetized with 0.5% proparacaine topical anesthesia. Animals were placed on a heating plate to keep their body temperature at 37°C. Responses were recorded using a gold loop wire electrode placed at the center of the cornea, a reference electrode in the mouth, and a ground electrode at the tail. A 2.5% hypromellose ophthalmic demulcent solution was used for corneal hydration. Dark-adapted ERG was performed using flashes with intensities ranging from 0.0001 to 10 sc cd.s/m^2^. Light adaptation was performed with white light at 20 sc cd/m^2^ for 2 minutes. Photopic ERG response was recorded using flashes with intensities ranging from 0.3 to 100 sc cd.s/m^2^.

### Optomotor Response

Optomotor response (OMR) was measured using the Optodrum (Striatech). Optodrum recommended parameters were used for the experiment. Square wave gratings were used at a fixed 99.72% contrast and rotation speed at 12 °/s. The number of cycles/° varied by a staircase algorithm, and the rotation of the drum was both clockwise and counterclockwise. The final determination of mouse visual acuity was determined by 2 successful and 3 unsuccessful trials.

### Optical Coherence Tomography (OCT) imaging

Cross sections of mouse retinas were imaged and acquired using the Heidelberg Spectralis HRA + OCT system (Heidelberg Engineering). Mice were anesthetized with an intraperitoneal injection of ketamine (100 mg/kg) and xylazine (6 mg/kg). Pupils were dilated with 0.5% tropicamide and 2.5% phenylephrine and anesthetized with 0.5% proparacaine topical anesthesia. 2.5% Hypromellose ophthalmic demulcent solution and Systane Ultra were used for corneal hydration until eyes were imaged. OCT imaging for each eye was centered around the optic nerve head. IR + OCT volume scans were obtained using Spectralis automatic real-time mode (ART), acquiring 55 cross-sectional images that were 200 μm apart. The individual layer thickness was measured using an AI OCT segmentation tool developed by the NEI IT Bioteam. Briefly, B-scan images that cross the center of the optic nerve were exported as a .tiff image. The AI segmentation tool outlines 6 lines on the OCT, totaling 5 retinal layers. Values for each layer were calculated from a region 350 to 650 um away from the center of the optic nerve head on each side. This region was chosen to have a relatively flat retinal layer thickness. Total retinal thickness is the summation of each retinal layer value.

### Statistical Analysis

Statistical analyses performed on GraphPad Prism 9.4.1 (GraphPad Software) included Welch’s t-test and one-way analysis of variance (Brown-Forsythe and Welch ANOVA) with post-hoc Tukey test. Chi-squared and Fisher’s exact tests with an α-value threshold of 0.05 was used to determine statistical significance.

### Study Approval

All animal experiments were conducted in accordance with recommendations of the Guide for the Care and Use of Laboratory animals of the National Institutes of Health (Protocol #NEI-680).

### Patient consent

This study was conducted under institutional review board-approved protocols in accordance with the Declaration of Helsinki for the release of clinical information and family history. Informed consent was obtained after the explanation of the study’s risks and benefits.

### Ethics approval

This study followed the tenets of the Declaration of Helsinki and was approved by all local ethics committees involved.

## Author Contributions

JL and RBH wrote and prepared the manuscript; JL, BG, HQ, and YZ performed data analysis; JL, YH, CL, MH, BG, ANia, CBen, NM, YS, LD, PL, JLei, and CH performed experiments and collected data; RS, HD, LAH, PA, YL, ANem, JTay, SD, MK, IM, MS, JTen, ATM, JLD, BM, SH, AV, CES, EIT, CBla, RS, VU, AW, MM, GA, MS, and RBH collected clinical data; all authors read and approved the final manuscript.

## Supporting information

Supporting Data Values

## Acknowledgements

We would like to thank the participating patients and their families, as well as the healthcare professionals involved in their care. We would like to thank Megan Kopera and her team for helping maintain and oversee the mouse colony. We would also like to thank Rudy Richardson for providing the protocol for the NTE esterase activity assay, as well as recommending vendors for chemicals related to the experiment. We would like to thank Jee Min Kim for editing the figures for the manuscript. We would like to thank Zubair Ahmed, Sally Camper and Sua Myong for their advice on the manuscript. Lastly, we would like to thank Ellen Sidransky and Nahid Tayebi for their tissue homogenizer and advice on the manuscript. This work was supported by National Eye Institute intramural funds. SH and AV are supported by Save sight society NZ, Retina NZ, and the Ombler trust. GA is supported by the National Institute of Health Research Biomedical Research Centre (NIHR-BRC) at Moorfields Eye Hospital and UCL Institute of Ophthalmology, a Fight For Sight UK Early Career Investigator Award (5045/46) and NIHR-BRC at Great Ormond Street Hospital Institute for Child Health. LP is supported by NEI K08-EY032098 and the Research to Prevent Blindness Career Development Award.

## Notes

### Competing Interest Statement

The authors have declared no competing interest.

