## Supplemental Figures and Tables for "Neuropathy target esterase activity predicts retinopathy among *PNPLA6* disorders"

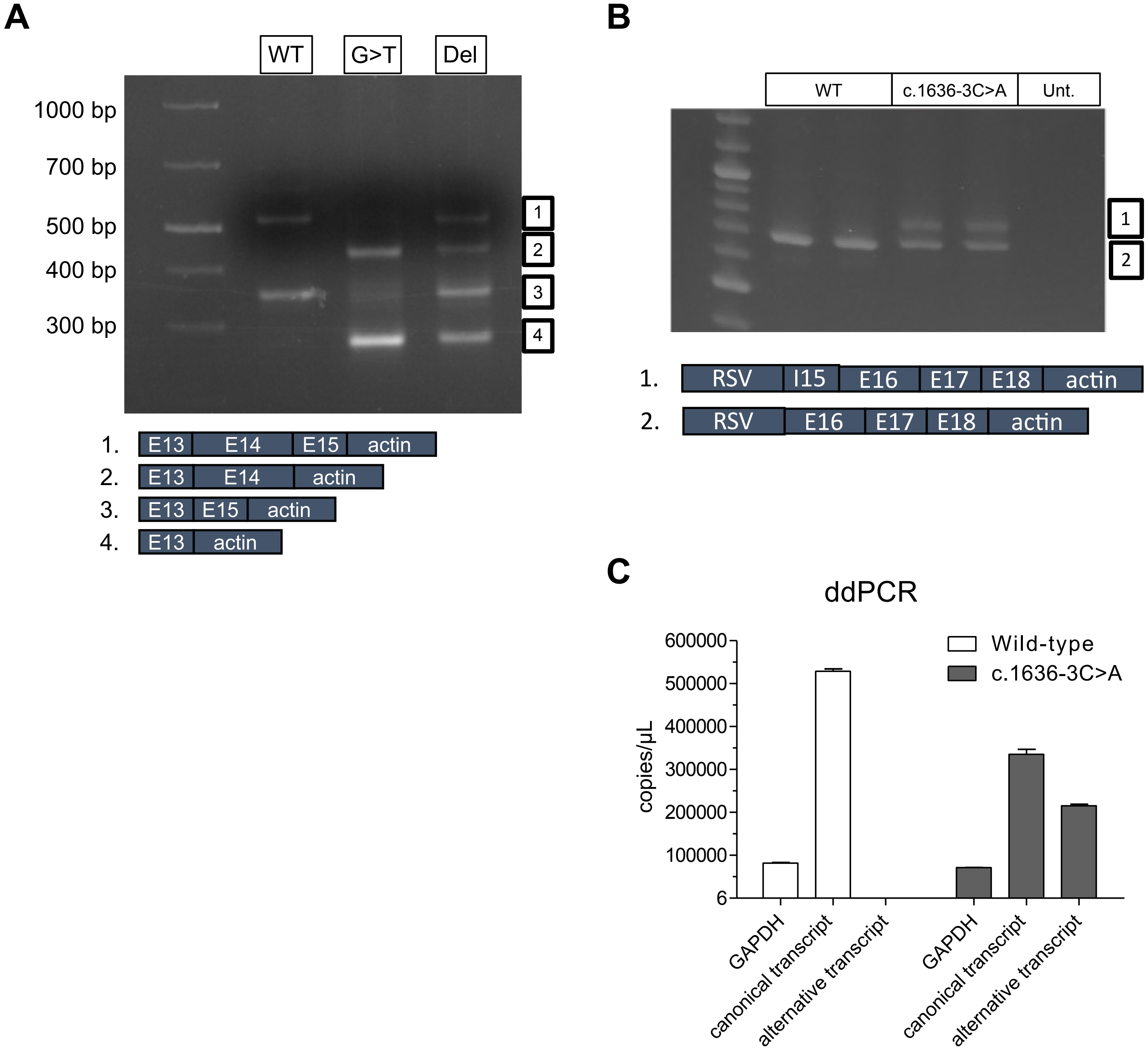


**Supplemental Figure 1. Minigene and ddPCR splicing assays of *PNPLA6* splicing variants. (A)** 3% Agarose gel electrophoresis of cDNA PCR products for *PNPLA6* WT, c.1635+3G>T (G>T), and c.1635+10_1635+15del (del) minigene assays. The WT lane has two bands, the top cDNA corresponds to Exons 13, 14, 15, and actin (~520bp) and the bottom band for Exons 13, 15, and actin (~350bp). Schematic of splice products as shown by Sanger sequencing of gel-purified cDNA PCR products. Numbers indicate location on the gel. **(B)** 2% Agarose gel electrophoresis of cDNA PCR products for *PNPLA6* WT (~685 bp), c.1636-3C>A(~685 + ~758 bp), and untransfected minigene assays, with schematic of the predicted splice products as shown by Sanger sequencing of gel-purified cDNA PCR products. Numbers indicate location on the gel. **(C)** ddPCR was used to quantify the percentage of alternative and canonical transcripts that were produced in the wild-type versus c.1636-3C>A variant.

**A**

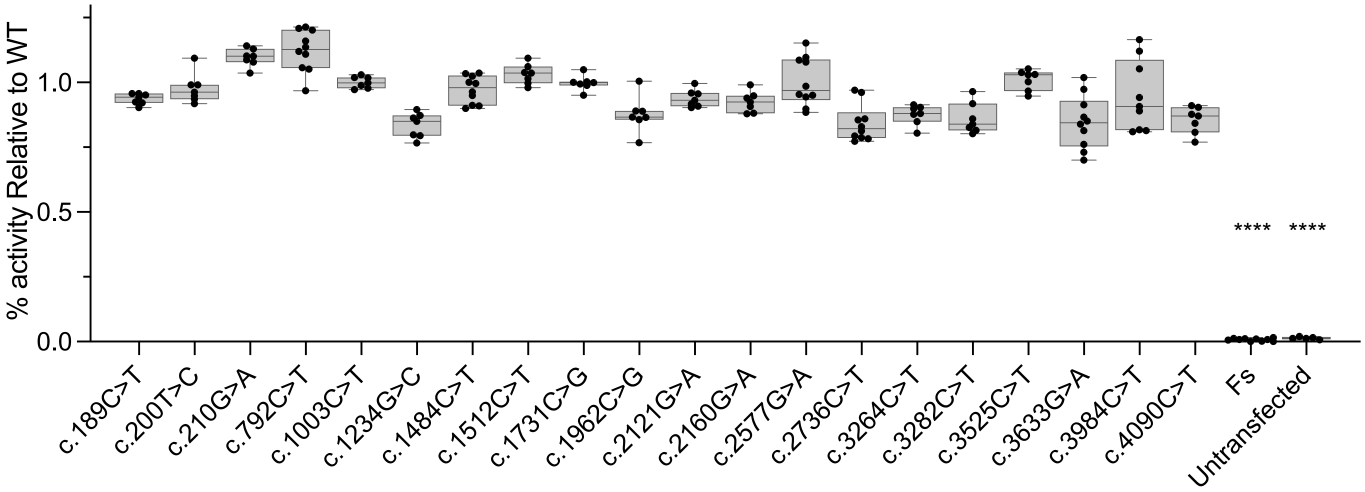


**Supplemental Figure 2. Activity of *PNPLA6* synonymous and likely benign variants.** Synonymous and missense variants were tested for residual hydrolase activity. Synonymous and missense benign variants were chosen based on an allele frequency in gnomAD >0.05%. N>7 for all samples tested. Each variant was not significantly different compared to the WT samples from the same run. Each run had a WT, a previously tested missense variant, Arg1031Glnfs*38, and Untransfected samples.

| Phenotype | NINDS-1 | OXF01_01 | Stanford CK | MS #1 | NEI_01 | Synofzik IHG26117 (2014A) | NEO_04 | Synofzik IHG25190 Sub. 1 (2014A) | Synofzik IHG25190 Sub. 2 (2014A) | Synofzik IHG25190 Sub. 3 (2014A) | Synofzik IHG25190 Sub. 4 (2014A) |
|---|---|---|---|---|---|---|---|---|---|---|---|
| Sex | M | F | F | M | M | M | F | F | M | F | F |
| Diagnosis | SPG39 | LNMS | SPG39 | BNHS | BNHS | SPG39 | OMCS | BNHS | BNHS | BNHS | BNHS |
| CNS | 3 | 17 | 35 | 7 | 11 | 4 | N/A | 6 | 6 | 7 | 7 |
| PNS | - | N/A | - | 8 | 11 | - | N/A | - | - | - | - |
| Endocrine | - | 16 | - | 4 | + | - | + | + | + | + | + |
| Ophthalmic | - | 9 | - | 2.5 | 6 | - | + | 2 | 3 | 1 | 2.5 |
| *PNPLA6* alleles | p.P1345S p.P1345S | p.G1129R p.T1305Hfs*6 | p.R1361* p.R1361* | p.R1099Q p.T1305Hfs*6 | p.L1051S p.L1051S | p.V1100G p.R1031Qfs*38 | p.L1051S p.P1056L | p.T1058I p.T1058I | p.T1058I p.T1058I | p.T1058I p.T1058I | p.T1058I p.T1058I |
| NTE Activity | 83% | 68% | 62% | 53% | 43% | 32% | 32% | 31% | 31% | 31% | 31% |

| Phenotype | Cyprus PA | Makuloluwa 2020 | NEI_02 | Hufnagel C:1 (2015) | Hufnagel D:1 (2015) | Hufnagel B:1 (2015) | Hufnagel E:1 (2015) | RS #1 | Cincinnati VU RS | Hufnagel A:1 (2015) |
|---|---|---|---|---|---|---|---|---|---|---|
| Sex | F | F | M | M | M | M | M | M | M | F |
| Diagnosis | OMCS | OMCS | OMCS | OMCS | OMCS | OMCS | OMCS | OMCS | OMCS | OMCS |
| CNS | - | - | 14 | - | - | - | 35 | N/A | - | - |
| PNS | - | - | - | - | - | 4 | 35 | N/A | - | - |
| Endocrine | 17 | 10 | 3 | + | 6 | 0.1 | 2 | + | 14 | 0.1 |
| Ophthalmic | 17 | 5 | 3 | + | 1 | 4 | 2 | 3 | 4 | 5 |
| *PNPLA6* alleles | p.A1112T p.R1183W | p.A1112T p.R1183W | p.T72fs* p.V1215A | c.1973+2T>G p.V1215A | dup(Ex14-20) p.V1215A | p.G1129R p.R1031Qfs*38 | p.G1129R p.R1031Qfs*38 | p.T1044A p.G1176S | p.R1183W p.R1031Qfs*38 | p.R1099Q p.G1176S |
| NTE Activity | 30% | 30% | 29% | 29% | 29% | 29% | 29% | 29% | 26% | 24% |

| Phenotype | Hufnagel A:2 (2015) | Synofzik IHG25357 (2014A) | NZ SH AV | NEI_03 | Cleveland ET | Invitae NY Z | Invitae NY R | POZ01_01 | Synofzik ARCA_05 Sub. 1 (2014A) | Synofzik ARCA_05 Sub. 2 (2014A) |
|---|---|---|---|---|---|---|---|---|---|---|
| Sex | M | M | F | M | F | M | M | M | F | F |
| Diagnosis | OMCS | BNHS | BNHS | OMCS | OMCS | OMCS | OMCS | OMCS | BNHS | BNHS |
| CNS | - | 20 | - | - | N/A | 5 | + | 1 | 27 | 6 |
| PNS | - | - | - | - | N/A | - | - | N/A | + | + |
| Endocrine | 0.1 | 14 | + | 0.1 | + | 3 | 14 | 1 | 14 | + |
| Ophthalmic | 4 | 14 | 4 | 10 | + | 3 | 1 | 2 | 12 | - |
| *PNPLA6* alleles | p.R1099Q p.G1176S | p.S1045L p.P1122L | p.R1147C p.Y496Rfs*19 | p.T1044A p.T1115M | p.S1045L p.P1126_D1128del | p.Y1109C p.R1031Qfs*38 | p.Y1109C p.R1031Qfs*38 | p.Q472* p.D1125N | p.V1110M p.V738Qfs*98 | p.V1110M p.V738Qfs*98 |
| NTE Activity | 24% | 24% | 22% | 20% | 18% | 17% | 17% | 12% | 11% | 11% |

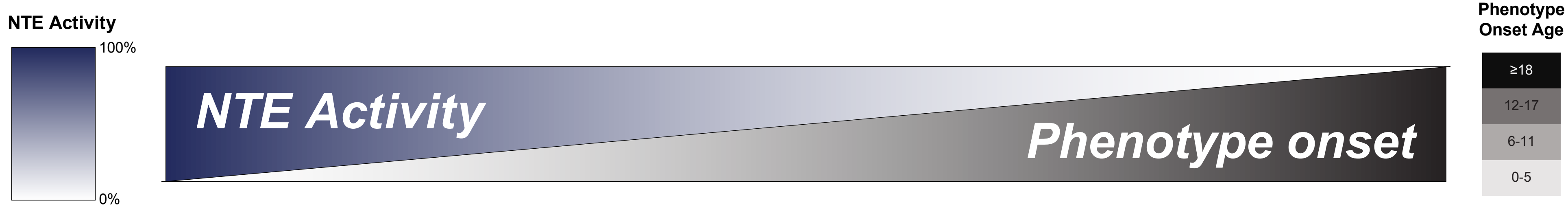


**Supplemental Figure 3. Clinical phenotype and genotype summary of unreported and reported patients in this study.** 23 unreported patients from our study cohort and 13 previously reported patients with biallelic *PNPLA6* variants (1, 4, 28).

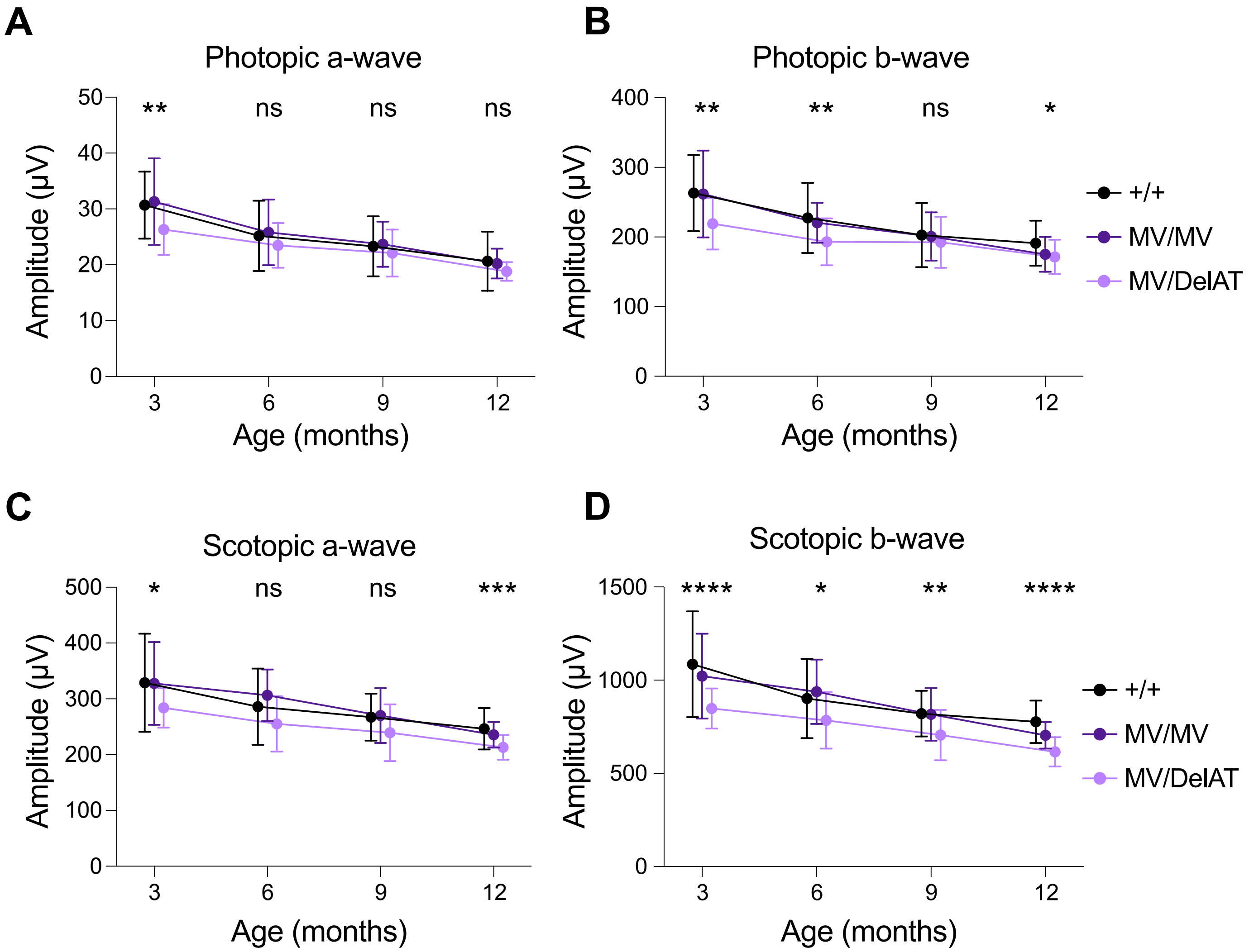


**Supplemental Figure 4. ERG time course of allelic series mice. (A-D)** ERG wave amplitudes at a stimulus intensity of 100 cd.s/m$^2$ (Photopic) and 10 cd.s/m$^2$ (Scotopic) for control, MV/MV, and MV/delAT mice under photopic or scotopic conditions over the course of 12 months. Control mice were littermates. Error bars indicate S.D. Significant values indicate significance between Control and MV/delAT values. Control and MV/MV line were not significantly different at all timepoints (exceptions include scotopic b-wave at 12 months). MV/MV and MV/delAT amplitudes were not significantly different (except for scotopic a-wave at 6 months, b-wave at 12 months, and photopic b-wave at 6 months). Each condition at each timepoint had an N=8-20. N values are found in Supporting Data Values xls.

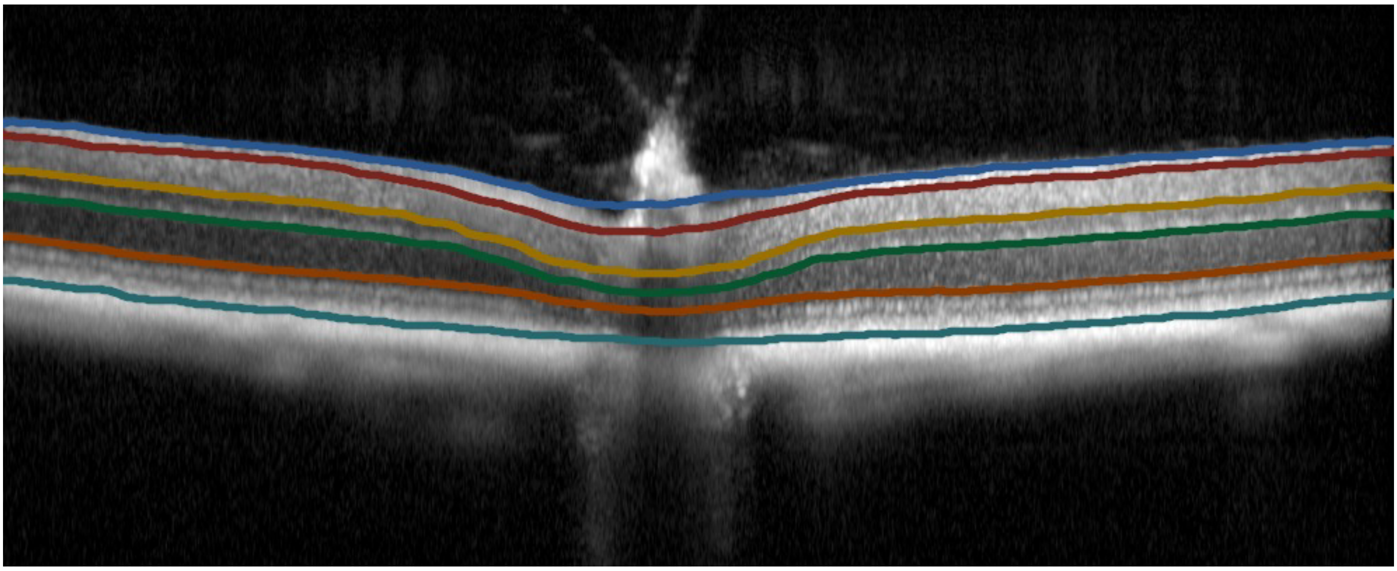

**Supplemental Figure 5. AI OCT segmentation tool validation.** Exported B-scan images that cross the center of the optic nerve. The AI segmentation tool outlines 6 lines on the OCT, totaling 5 retinal layers (blue to red: nerve fiber layer; red to yellow: inner plexiform layer; yellow to green: inner nuclear layer; green to orange: outer nuclear layer; orange to teal: photoreceptor and RPE).

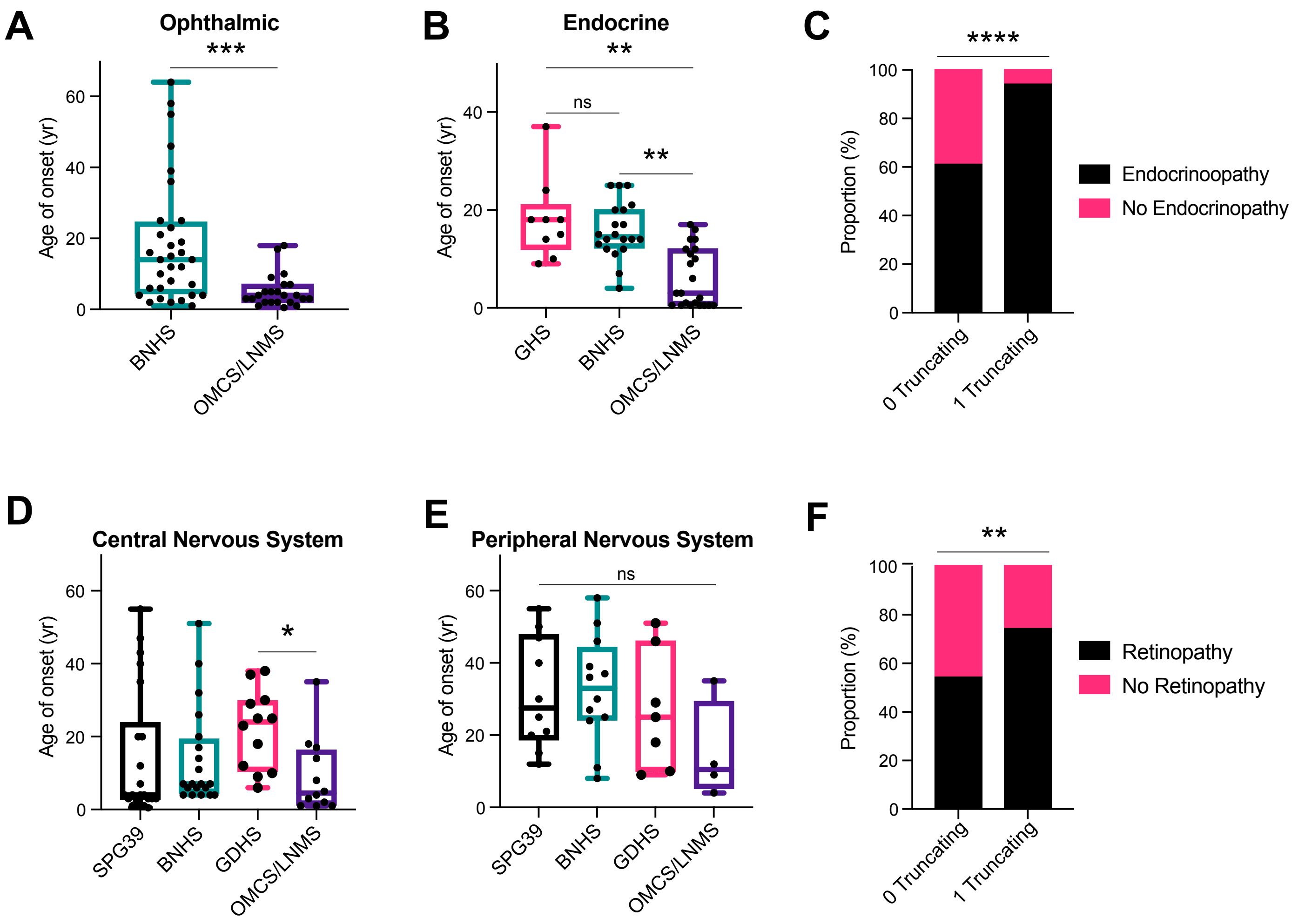


**Supplemental Figure 6. Affected tissue age of onset in patients with *PNPLA6* variants is significantly different in ophthalmic and endocrine tissue, but not nervous system tissue. (A)** Comparing ophthalmic tissue age of onset between BNHS and OMS/LMS patients. **(B)** Comparing endocrine tissue age of onset between GDHS, BNHS, and OMS/LMS patients. **(C)** Frequency distribution of patients with and without endocrinopathy, grouped by 0 vs 1 truncating genotype. **(D)** Comparing CNS tissue age of onset across all syndromes. **(E)** Comparing PNS tissue age of onset across all syndromes. **(F)** Frequency distribution of patients with and without retinopathy, grouped by 0 vs 1 truncating genotype. (B), (D), (E) used a Brown-Forsythe ANOVA with post-hoc Tukey test. A) used Welch's t-test with α=0.05. p > 0.05 = ns, p < 0.05 = *, p < 0.01 = **, p < 0.001 = ***, p < 0.0001 = ****. (C) and (F) used a Fisher's exact test with α =0.05. Error bars indicate S.D. N values found in Supporting Data Values xls.

| Genotype | M1030V | V1185A | R1069Q | G1146S | delAT |
|---|---|---|---|---|---|
| **M1030V** | **VIABLE** (14/49) | **VIABLE** (12/33) | **VIABLE** (11/43) | **VIABLE** (5/59) | **VIABLE** (15/66) |
| **V1185A** | | **VIABLE** (11/60) | Not Viable (0/52) | Not Viable (0/50) | Not Viable (0/81) |
| **R1069Q** | | | Not viable (0/56) | Not Viable (0/42) | Not Viable (0/59) |
| **G1146S** | | | | Not Viable (0/61) | Not Viable (0/49) |
| **delAT** | | | | | Not Viable (0/30) |

**Supplemental Table 1. Viability Matrix of *Pnpla6* allelic series.**
